# Spontaneously emerging patterns in human visual cortex and their functional connectivity are linked to the patterns evoked by visual stimuli

**DOI:** 10.1101/518712

**Authors:** DoHyun Kim, Tomer Livne, Nicholas V. Metcalf, Maurizio Corbetta, Gordon L. Shulman

## Abstract

The function of spontaneous brain activity is an important issue in neuroscience. Here we test the hypothesis that patterns of spontaneous activity code representational patterns evoked by stimuli and tasks. We compared in human visual cortex multi-vertex patterns of spontaneous activity to patterns evoked by ecological visual stimuli (faces, bodies, scenes) and low-level visual features (e.g. phase-scrambled faces). Specifically, we identified regions that preferred particular stimulus categories during localizer scans (e.g. extra-striate body area for bodies), measured multi-vertex patterns for each category during event-related task scans, and then correlated over vertices these stimulus-evoked patterns to the pattern measured on each frame of resting-state scans. The mean correlation coefficient was essentially zero for all regions/stimulus categories, indicating that resting multi-vertex patterns were not biased toward particular stimulus-evoked patterns. However, the spread of correlation coefficients between stimulus-evoked and resting patterns, i.e. both positive and negative, was significantly greater for the preferred stimulus category of an ROI (e.g. body category in body-preferring ROIs). The relationship between spontaneous and stimulus-evoked multi-vertex patterns also governed the temporal correlation or functional connectivity of patterns of spontaneous activity between individual regions (pattern-based functional connectivity). Resting patterns related to an object category fluctuated preferentially between ROIs preferring the same category, and patterns related to different categories fluctuated independently within their respective preferred ROIs (e.g. body- and scene-related multi-vertex patterns within body- and scene-preferring ROIs). These results support the general proposal that spontaneous multi-vertex activity patterns are linked to stimulus-evoked patterns, consistent with a representational function for spontaneous activity.

## Introduction

Spontaneous neural activity is observed throughout the brain, yet its function remains mysterious. An important clue, however, comes from work that has uncovered striking similarities between spontaneous activity and the activity evoked by a task (Arcaro et al 2015, Berkes et al 2011, Cole et al 2016, Fiser et al 2004, Fox et al 2007, Heinzle et al 2011, Kenet et al 2003, Omer et al 2018, Raemaekers et al 2014, Ryu & Lee 2018, Tavor et al 2016, Tsodyks et al 1999). For example, the temporal correlation of spontaneous activity between brain regions (functional connectivity, FC) closely resembles the spatial topography of task-evoked activity (Biswal et al 1995, de Pasquale et al 2010, Greicius et al 2003, He et al 2008, Power et al 2011, Yeo et al 2011), links distributed brain regions into functional networks, and can be used to predict task activation (Cole et al 2016, Osher et al 2019, Tavor et al 2016).

The remarkable spatiotemporal regularities of spontaneous activity and widespread findings that abnormalities in inter-regional correlations of spontaneous activity in humans are associated with neurological and psychiatric disorders (e.g. (Grefkes & Fink 2014, Northoff & Duncan 2016)) have motivated a search for functional explanations. One hypothesis is that spontaneous activity has a role in the synaptic homeostasis of structural connections (Deco et al 2013). Another idea is that fluctuations of spontaneous activity between regions constitute a spatiotemporal prior that facilitates the recruitment of task circuitries during behavior (Petersen & Sporns 2015, Raichle 2011). A third hypothesis is that spontaneous activity has a role in representing information about external (Fiser et al 2010) and internal states (Harmelech & Malach 2013). Genetically determined circuitries generate spontaneous activity that is shaped in the course of development by experience through Hebbian statistical learning (Berkes et al 2011). Conversely, spatial and temporal patterns of spontaneous activity constrain task-evoked patterns. As a result of this cyclic process, both spontaneous and task-evoked activity code similar representations of internal and external states (Fiser et al 2010). The same process determines the spontaneous interactions between regions, which reflect connectivity patterns that are coded as synaptic efficacies in cortical networks (Harmelech & Malach 2013, Strappini et al 2018). Figure 1 illustrates schematically the representation hypothesis of spontaneous brain activity.

**Figure 1.**
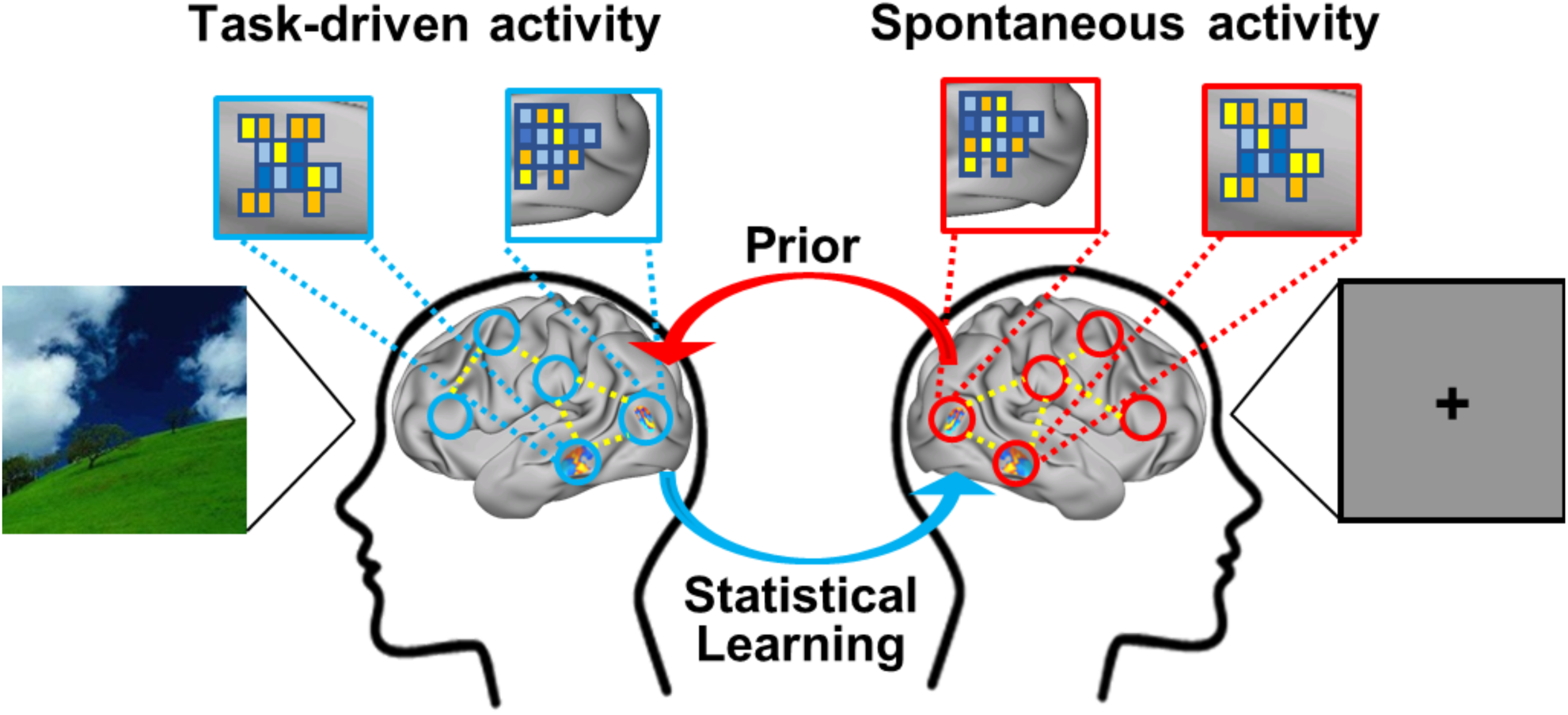
The putative cyclic interplay between brain activity evoked by real-world experiences and resting-state activity.

The *representation* hypothesis has been supported by human and animal work. In animals, imaging of neural activity at a scale that extends across many cortical columns has shown that the macro-scale spatial pattern of spontaneous neural activity within a sensory area in an anesthetized animal mirrors the pattern of activity evoked by stimulation of a specific visual feature (Kenet et al 2003, Omer et al 2018). In humans, fMRI studies of early visual cortex have shown that FC between individual voxels respects the stimulus-evoked selectivity of voxels for polar angle, eccentricity, and low-level stimulus features (Arcaro et al 2015, Heinzle et al 2011, Raemaekers et al 2014, Ryu & Lee 2018). Recent work has also shown that voxel-wise resting FC in visual cortex is better approximated by the FC evoked by movies than by more artificial stimuli such as rotating checkerboards or static pictures of stimuli (Strappini et al 2018, Wilf et al 2017). Finally, Chen and colleagues (Chen et al 2017) measured the voxelwise resting FC between regions of ventral visual cortex and a second region that was functionally related to a specific visual category (e.g. tools). The multivoxel pattern of voxelwise FC values in a ventral visual region was similar to the multivoxel pattern of activity evoked in that region by exemplars of the category.

Here we further test the representation hypothesis in human visual cortex by determining whether multi-vertex patterns of resting activity resemble stimulus-evoked patterns of activity. This analysis bears some resemblance to a representational similarity analysis, which is a standard methodology for identifying the representational content of evoked activity in a brain region (Kriegeskorte et al 2008). Unlike Chen et al (Chen et al 2017), we examine the multi-vertex pattern of resting activity at each timepoint rather than separately analyzing each individual vertex over timepoints in a resting FC analysis.

Specifically, we measured the similarity of resting and stimulus-evoked multi-vertex patterns in different category specific regions of human visual cortex (e.g. extrastriate body area, EBA, fusiform face area, FFA, and parahippocampal place area, PPA), for different stimulus categories (e.g. bodies, faces, places). We also examined the similarity of resting and stimulus-evoked patterns in early visual cortex and in category specific regions of visual cortex for the low-level features comprising phase-scrambled and grid-scrambled stimuli. Finally, we examined whether in the resting-state, fluctuations over time in the amplitude of stimulus-evoked patterns were correlated across regions (hereafter termed ‘pattern-based functional connectivity’). For example, do fluctuations in the amplitude of a resting multi-vertex pattern of activity in a region that is related to a particular category (e.g. bodies) correlate over time with fluctuations in the amplitude of the multi-vertex pattern of activity for the same category in a different region? Because this analysis measures the correlated fluctuations of spontaneous patterns of activity across regions, it differs from a standard functional connectivity (FC) analysis, in which the activity of a single voxel is temporally correlated with the activity of other voxels, or the activity averaged across the voxels of a region is temporally correlated with the activity of other voxels or averaged regions. Similar multi-vertex, pattern-based FC analyses have previously been reported, but only for task-evoked activity, not for resting activity (Anzellotti & Coutanche 2018, Chen et al 2018, Coutanche & Thompson-Schill 2013).

The representation hypothesis of spontaneous activity makes several specific predictions about the expected multi-vertex pattern of activity within a region and the interaction of that pattern with the patterns from other regions. First, if spontaneous activity carries information about stimulus categories such as faces, bodies, or scenes, then spontaneous multi-vertex activity patterns, i.e. patterns of activity observed at rest in the absence of any stimulation, in functionally specialized occipital regions such as the extrastriate body area (EBA, (Downing et al 2001)) should be more related to the activity patterns evoked by a preferred stimulus category (e.g. bodies) than by non-preferred categories. This predicted relationship between spontaneous and task-evoked multi-vertex patterns putatively reflects the entrainment of task-evoked patterns into spontaneous activity during development and through experience. Second, if the FC between regions reflects in part correlated fluctuations of the spontaneous representational content of those regions, then the magnitude of resting pattern-based FC between two regions should depend on whether the regions prefer the same visual stimulus category and on whether the tested patterns that are being correlated in the two regions code that preferred category.

## Materials and Methods

### Participants

The study included 16 healthy young adult volunteers (10 female; age 21 – 35 years-old) with no prior history of neurological or psychiatric disorders. All participants were right-handed native English speakers with normal or corrected-to-normal vision. All participants gave informed consent to take part in the experiment, and the study was approved by the Institutional Review Board (IRB) of Washington University in St. Louis School of Medicine.

### Stimuli

Nine categories of color images subtending 8° x 8° of visual angle were included in event-related ‘task’ fMRI scans. Seven ‘whole-object’ categories consisted of images that are encountered in real-world environments: human faces, human bodies, mammals, chairs, tools, scenes, and words. Stimuli, excluding the word category, were obtained from Downing et al., 2006 (Downing et al 2006). Faces, bodies and mammals served as animate categories, and chairs, tools and scenes as inanimate categories (Kriegeskorte et al 2008). Word stimuli were included for exploratory analyses and results for those stimuli will not be considered in this paper.

Two control stimulus categories were constructed from the above stimuli excluding the word stimuli. A low-level control consisted of phase-scrambled stimuli that preserved the spatial frequency amplitude spectrum of the whole-objects images. An intermediate-level control consisted of grid-scrambled stimuli that included basic visual properties of the whole-objects images such as line segments and connectors. For the low-level control condition, 2D phase-scrambled images of the exemplars from the 6 categories were generated by applying the same set of random phases to each 2-dimensional frequency component of the original image while keeping the magnitude constant (Watson et al 2016). Exemplars from all six whole-objects categories except real word stimuli were 2D phase-scrambled, yielding a total of 144 2D Phase-scrambled stimuli. For the intermediate-level control condition, grid-scrambled images of exemplars from the six whole-objects categories were generated by sub-dividing each image into a 10 x 10 grid (each grid is 0.8° x 0.8°) and randomly rearranging the individual grid segments. Figure 2 shows exemplar stimuli from the 8 categories used.

**Figure 2.**
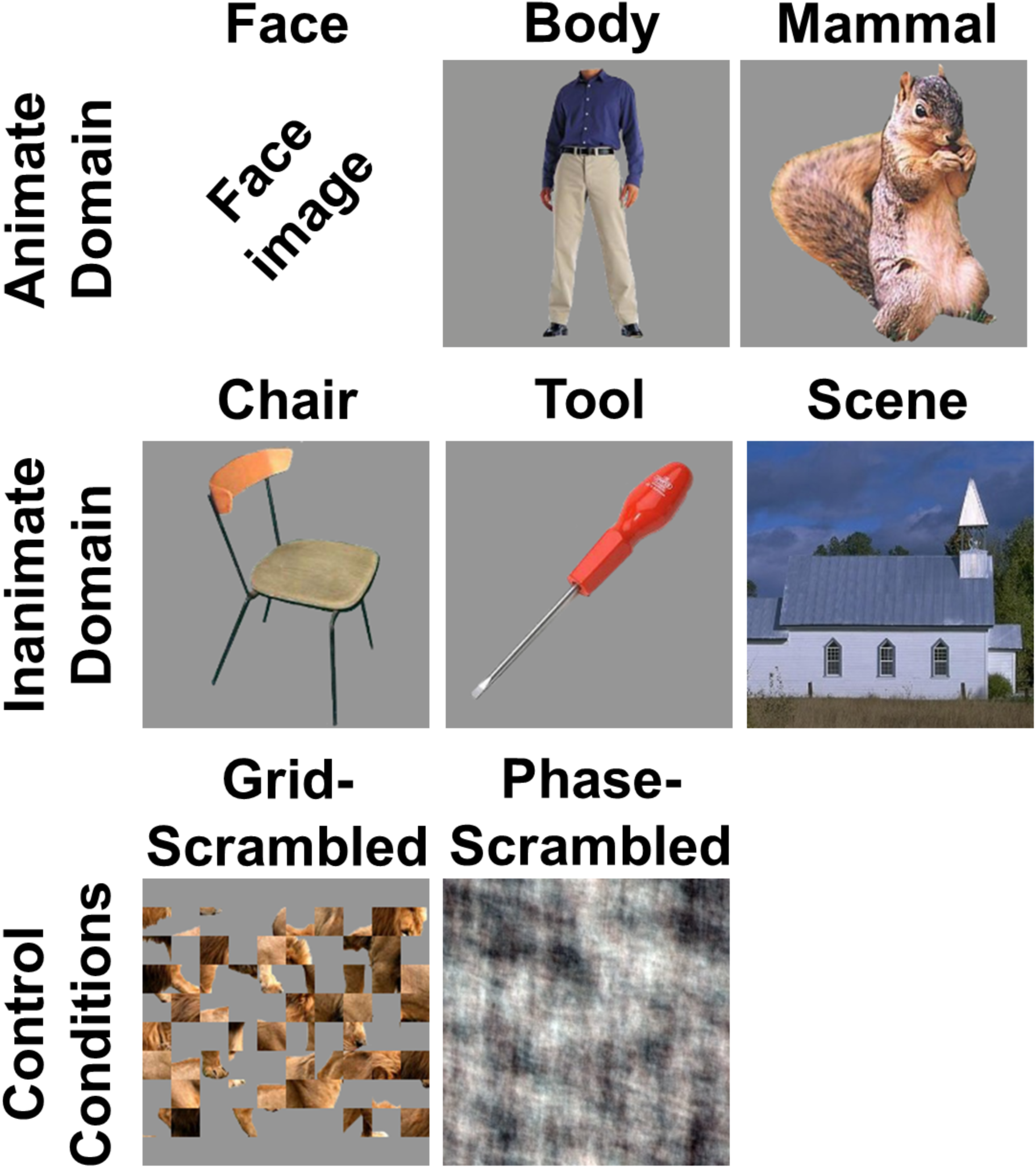
Stimulus categories used in the experiment.

Color images of exemplars from seven categories were included in blocked-design localizer scans: human faces, human bodies, objects (chairs and tools), scenes, words, false font character strings and phase-scrambled images. The categories for the localizer scans differed slightly from the categories for the task scans since the former was only used to define the regions of interest (ROIs). ROIs related to the false font and word stimuli will not be considered in this paper.

Stimuli were presented using the Psychophysics Toolbox package (Brainard 1997) in MATLAB (The MathWorks). Stimulus images were projected onto a screen and were viewed through a mirror mounted on the head coil. All stimuli were presented centrally on a gray background.

### Scanning Procedure

The study consisted of two separate sessions, each conducted on a separate day (Fig. 3). In session one, subjects received 3 resting state runs, 2 localizer runs, and 8 task runs. In session 2, subjects received 2 resting state runs, 2 localizer runs, 8 task runs, and 2 post-task resting state runs. One subject had a total of 13 task runs over the two sessions instead of 16.

**Figure 3.**
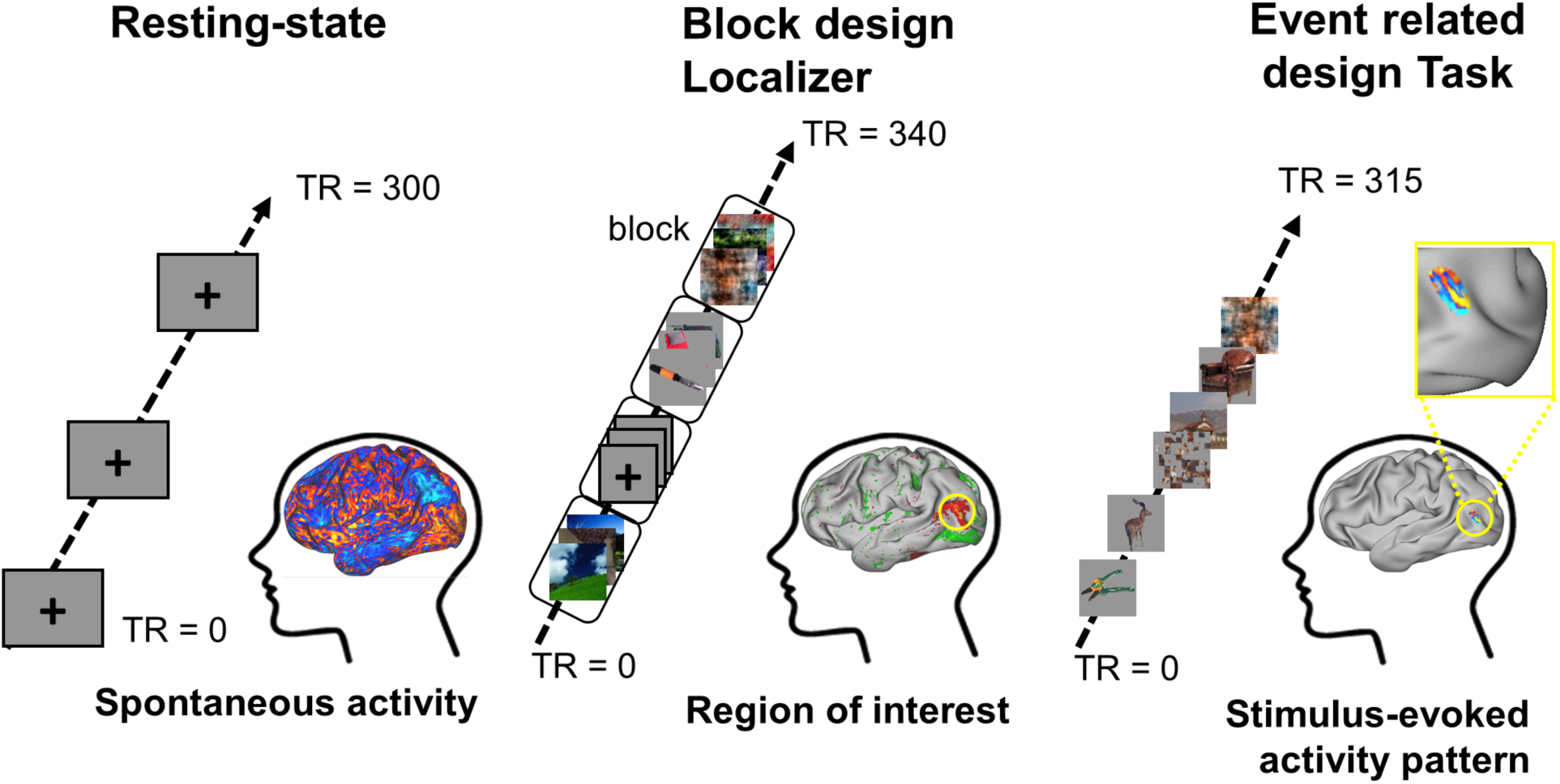
Experimental design, with separate resting scans, blocked-design localizer scans, and event-related task scans.

#### Resting state runs

Participants received a total of 7 resting state scans, each lasting 5 min (300 TRs). During a scan the participant was asked to maintain fixation on a cross that was displayed at the center of the screen during the entire run. Five resting scans (3 for the first session and 2 for the second session) were conducted before any localizer or task scans to collect stimulus-free intrinsic activities. For the second session only, two additional 5 min resting state scans were conducted after the task scans to investigate potential post stimuli-driven effects on intrinsic activity. The results from the post-task resting scans will not be discussed here.

#### Localizer runs

Each session included 2 localizer runs (4 in total), each lasting 5 min and 40s (340 TRs), and each localizer scan was presented in a blocked fMRI design. Each block of a localizer run contained 20 images of a single category, and those images were different from the images used in the task scans. A fully randomized sequence of eight blocks, consisting of the 7 stimulus categories and a fixation block, was repeated twice within each run. At the beginning and the end of each run, an additional fixation block was presented for 4s and 16s. Within each category block, images were presented for 300ms with an inter-stimulus interval (ISI) of 700ms. A fixation cross was continuously present at the center of the screen during the ISI and during fixation blocks. During category blocks, participants performed a minimally cognitively engaging task by pressing a button if the initially presented image was changed in size or position during the 300ms presentation.

#### Task runs

Each session included 8 task runs (16 in total), each lasting 5 min and 15s (315 TRs). For each subject and for each run, stimulus presentation order and inter-stimulus intervals were fully randomized using Optseq2 (Dale 1999). Each stimulus presentation lasted for 300ms and the interval between stimuli was jittered between 3.7s and 8.7s. A fixation cross was continuously present at the center of the screen during the ISI. In each whole-objects category, there were 24 separate exemplars (e.g. 24 different faces) and each exemplar was repeated 4 times. In each scrambled category, there were 96 exemplars, each presented once. Participants performed a minimally cognitively engaging task by pressing a button if the presented image changed its size or position during a 300ms presentation, the same task as that performed during the localizer scans.

### Imaging Parameters and fMRI pre-processing

Structural and fMRI images were obtained from a Siemens 3T Prisma MRI scanner. Structural images for atlas transformation and lesion segmentation were acquired using T1-weighted magnetization prepared-rapid gradient echo (MP-RAGE) (1 x 1 x 1 mm voxels; echo time [TE] = 2.36 ms, repetition time [TR] = 1700 ms, TI=1000 ms, flip angle = 8°) and T2-weighted fast spin echo sequences (1 x 1 x 1 mm voxels; TE = 564 ms, TR = 3200 ms). FMRI scans were collected using a gradient echo-planar sequence sensitive to BOLD contrast (TE = 26.6 ms, flip angle = 58°, 2.4 x 2.4 x 2.4 mm voxels, 48 contiguous slices, TR = 1.0 s, and multiband factor of 4).

### fMRI pre-processing

fMRI data underwent pre-processing as previously described (Siegel et al 2016). This included: (1) compensation for asynchronous slice acquisition using sinc interpolation; (2) elimination of odd/even slice intensity differences resulting from interleaved acquisition; (3) whole brain intensity normalization to achieve a mode value of 1000; (4) spatial realignment within and across fMRI runs; and (5) resampling to 2.4 mm cubic voxels in atlas space, including realignment and atlas transformation in one resampling step. Cross-modal (e.g. T2-weighted to T1-weighted) image registration was accomplished by aligning image gradients.

Surface generation and processing of functional data followed procedures similar to Glasser et al (Glasser et al 2013). First, anatomical surfaces were generated for each subject’s T1 MRI using FreeSurfer automated segmentation (Fischl et al 1999). This step included brain extraction, segmentation, generation of white matter and pial surface, inflation of the surfaces to a sphere, and surface shape-based spherical registration to the subjects’ “native” surface to the fs_average surface. The left and right hemispheres were then resampled to 164,000 vertices and registered to each other (Van Essen et al).

Data were passed through several additional preprocessing steps: (i) removal by regression of the following sources of spurious variance: (a) six parameters obtained by rigid body correction of head motion, (b) the signal averaged over the whole brain (global signal regression), (c) signal from ventricles and CSF, and (d) signal from white matter; (ii) temporal filtering retaining frequencies in the 0.009–0.08-Hz band; and (iii) frame censoring (framewise displacement (FD) ≥ 0.5mm). The first four frames of each BOLD run were excluded.

To account for magnitude variability between different task and resting state runs, the BOLD timeseries for each vertex were Z-normalized across time within the task and the resting state runs. This Z-normalization was not applied to the localizer scans. Also, it was not applied to the Task scans for a separate analysis described below in which task-evoked activation magnitudes were determined (see below, ***Task scans: multi-vertex activation patterns)*.**

### Defining ROIs from localizer activation contrasts

The next step was to define ROIs for each subject that showed a preference for specific categories or features. For this purpose, subject-specific ROIs were defined from univariate vertex-wise statistical contrasts on the localizer activation magnitudes for different categories. For example, face-selective areas were defined from the vertices for the contrast of faces minus objects, where objects consisted of chairs and tools. First, for each participant a general linear model (GLM) was applied to their functional localizer scans. The GLM consisted of separate regressors for each stimulus category (e.g. faces) using an assumed hemodynamic response function from the Statistical Parametric Mapping (SPM12), a baseline term, and a linear trend term. Condition contrasts were formed to identify vertices showing a preference for each category, using a scheme similar to that of Bracci and Op de Beeck (2016) (Bracci & Op de Beeck 2016): body-preference (body > objects, i.e. chairs and tools), face-preference (face > objects), scene-preference (scene > objects), whole-objects-preference (face + body + scene + object (chair+tool) > phase-scrambled), and phase-scrambled-objects-preference (phase-scrambled > face + body + scene + object (chair+tool)).

A group random-effect statistical Z-map for each contrast was then computed from the single-subject localizer GLMs (see Fig. 4A for the group z-statistic maps for body-, face-, and scene-preferences). The Z-values obtained were sorted in magnitude. From the highest Z-values from the map, the group peak with the next highest Z-value was generated until the Z-value was <= 2.0. Group peaks had to be separated by at least 38.4mm (9.6 mm x 4) in the sphere mesh to prevent a vertex being assigned to multiple ROIs in a subject. ROIs were then defined separately for each participant based on the individual’s univariate statistical maps (Oosterhof et al 2012, Wurm et al 2016). From each group peak defined above, the corresponding peak for an individual subject peak was defined as the vertex with the highest Z-value within a sphere of 9.6 mm radius centered around the group peak in each subject’s sphere mesh. The single-subject ROI was formed from the vertices exceeding Z= 2.0 in a sphere of 9.6 mm radius centered around the peak in the subject’s mesh. All ROIs used in the following analysis contained at least 175 vertices in at least 14 subjects. ROIs in individual subjects with less than 175 vertices were discarded.

**Figure 4.**
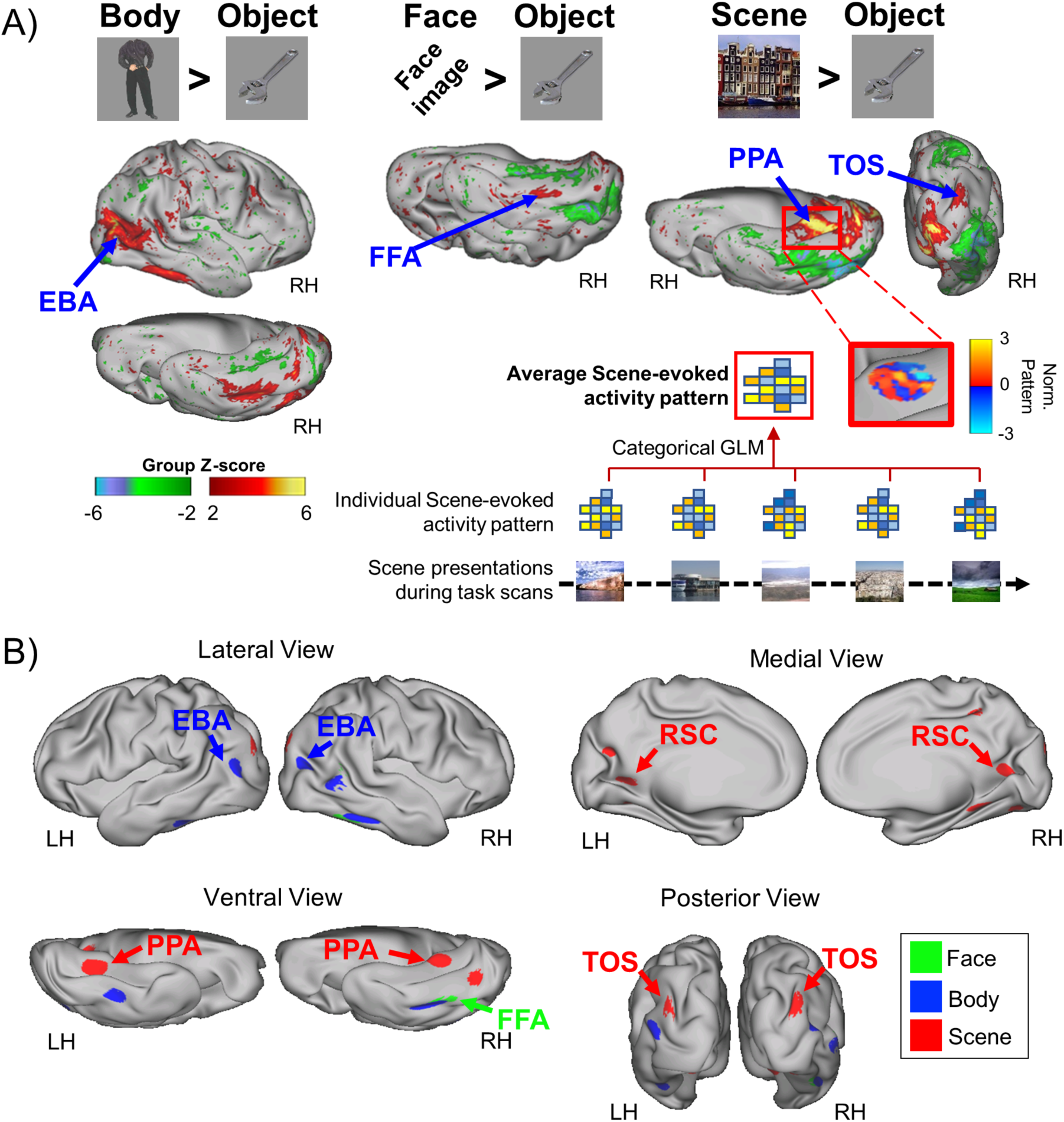
Body, Face, and Scene ROIs (regions of interest). **(A)** Group z-statistic Localizer maps for category-preferring visual regions. ROIs were separately defined for each individual from their localizer maps using the group foci as a constraint. **(B)** Schematic rendering of three sets of category-preferential ROIs for faces, bodies, and scenes using the object category (tools and chairs) as the baseline.

To remove differences in BOLD magnitude across MR frames, for each ROI a z-normalization was applied across the vertices of each frame of the resting and task scans. This within-frame z-normalization was not applied to the localizer scans. Also, it was not applied to the Task scans for a separate analysis described below in which task-evoked activation magnitudes were determined (see below, ***Task scans: estimation of multi-vertex activation patterns)*.**

Two sets of ROIs were created for use in different analyses. The first set was created from the localizer-defined ROIs that preferred a specific category (face, body, or scene) as compared to the object category (chairs + tools), and was used to compare the similarity of stimulus-evoked and resting multi-vertex patterns in high-level, category-preferring regions of visual cortex. Vertices from all ‘constituent’ ROIs that preferred a category (e.g. bodies) were grouped into a single ‘joint-ROI’, excluding all vertices located in early visual areas (V1 to V3) (Strappini et al 2018), as estimated from surface topology using the template created by Wang et al. (Wang et al 2015). For instance, the body joint-ROI included constituent regions such as left and right EBA, left and right fusiform body area (FBA), and the scene joint-ROI included constituent regions such as PPA, the transverse occipital sulcus (TOS), and retrosplenial cortex (RSC).

The rationale for combining all the vertices that prefer a category into a single ‘joint’ ROI was as follows. First, we had no apriori reason in our analysis of spontaneous activity patterns to expect different results for regions preferring the same category. Second, the use of joint-ROIs reduced the total number of statistical comparisons in our subsequent analyses of spontaneous activity patterns, simplifying the analysis. Third, the use of joint-ROIs increased the number of vertices over which spontaneous activity patterns were assessed, increasing the reliability of the analysis.

A second set of contrasts identified ROIs that preferred whole-objects relative to phase-scrambled objects (face + body + scene + object > phase-scrambled) or the reverse (phase-scrambled > face + body + scene + object). This set of ROIs was used to compare resting and task-evoked patterns for whole objects vs. low level features. Whole-Object and Phase-Scrambled Object ‘constituent’ ROIs were grouped, respectively, into a Whole-Objects joint-ROI and a Phase-Scrambled Objects joint-ROI. Table 1 summarizes the mean MNI coordinate, mean Z-score for the obtained group peak, and mean number of vertices for all constituent ROIs in each joint-ROI. Figure 4B schematically indicates the position of all constituent ROIs in the Face-, the Body- and Scene-joint-ROIs based on their group peak locations. Figure 5B shows the location of all constituent ROIs in the Whole-Objects joint-ROI and the Phase-Scrambled Objects joint-ROI superimposed on a surface map of V1-V3 using the template from Wang et al. (Wang et al 2015).

**Figure 5.**
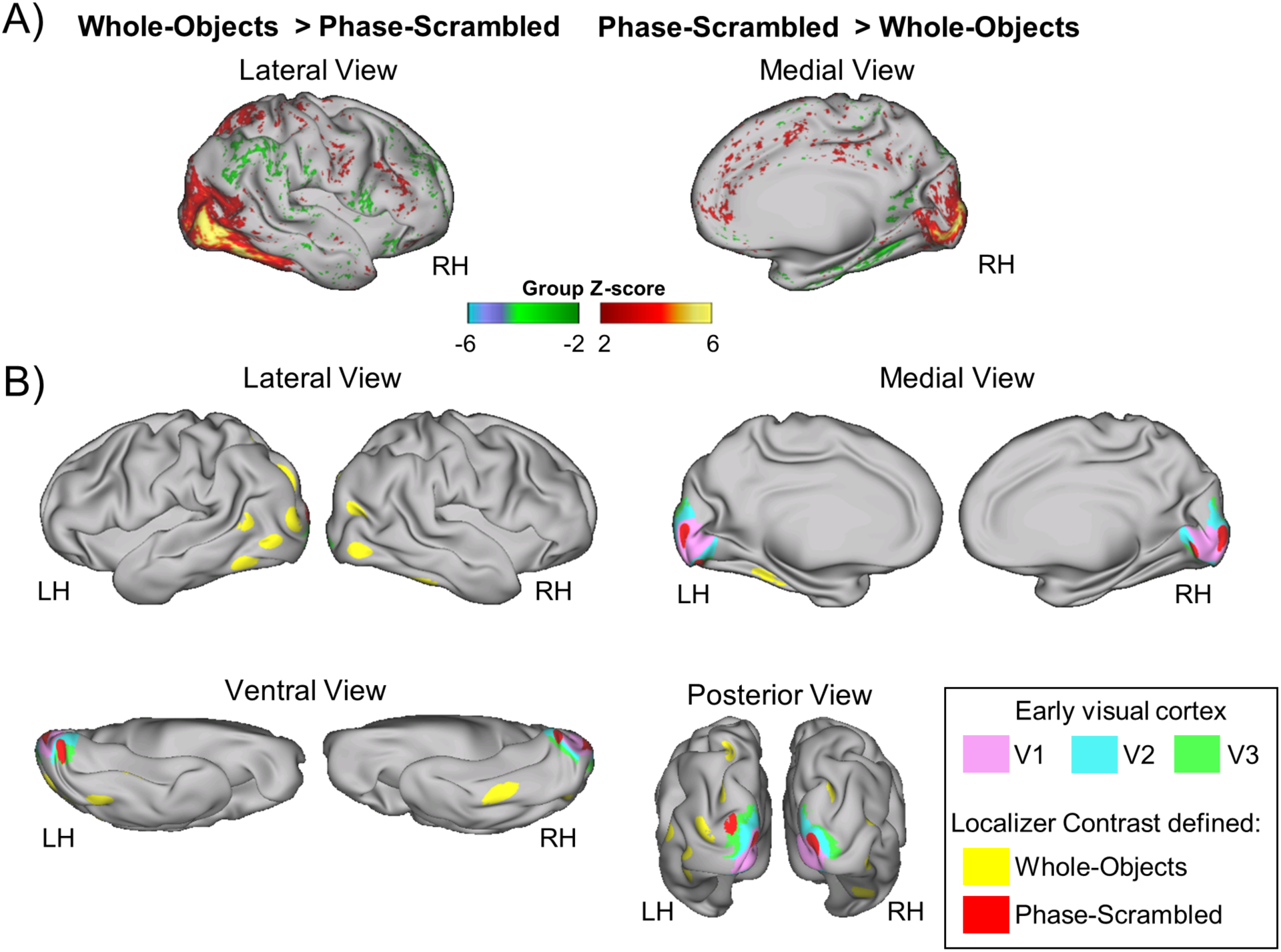
Whole-object and Phase-scrambled ROIs. **(A)** Group z-statistic Localizer maps of visual regions that prefer whole-objects or phase-scrambled objects. **(B)** Schematic rendering of Whole-Objects and Phase-Scrambled ROIs. ROIs were separately defined for each individual from their localizer maps using the group foci as a constraint. Surface renderings of V1, V2, and V3 from the Wang et al. template (Wang et al 2015) are superimposed.

**Table 1.**
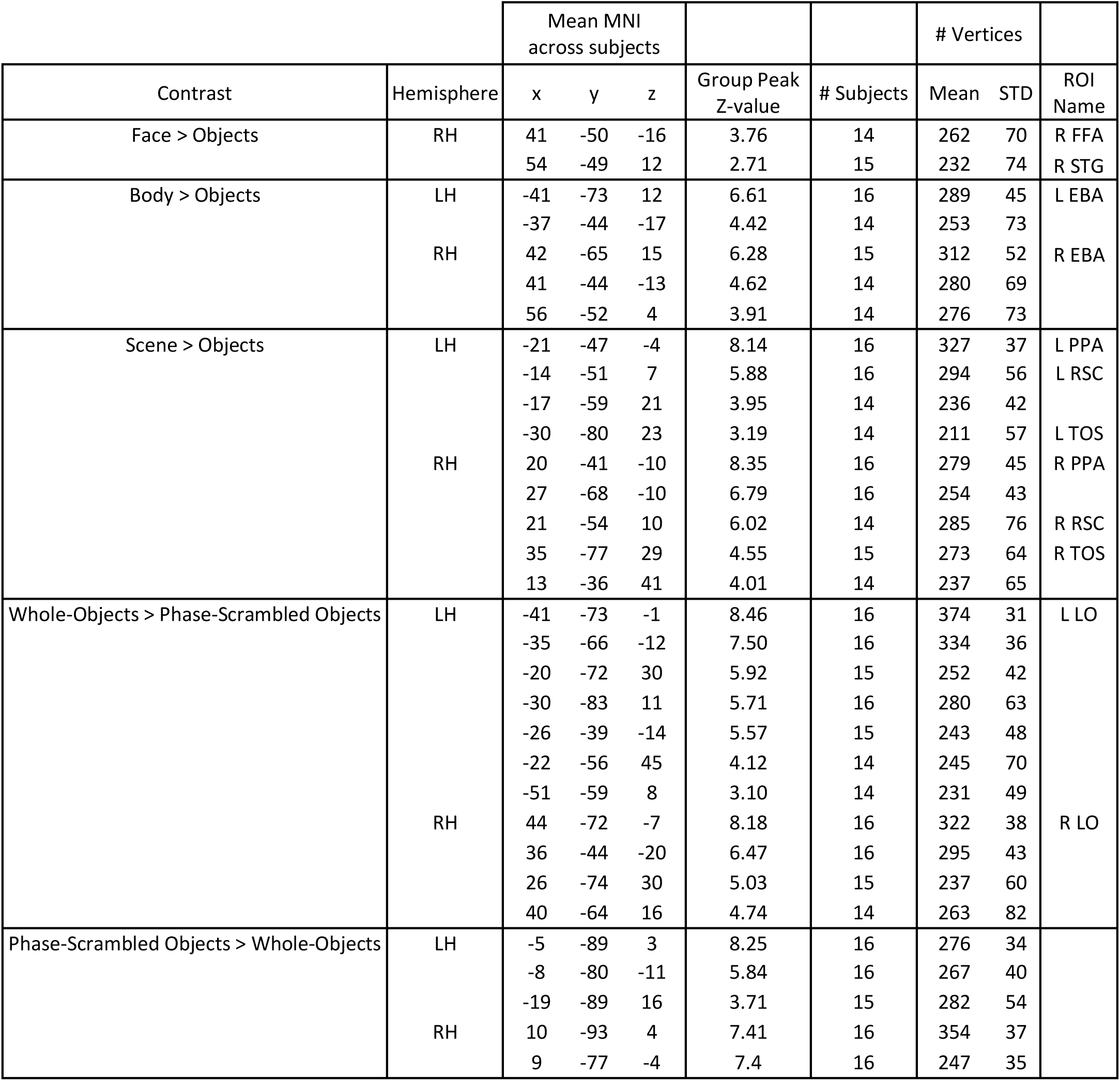
ROI Summary

### Task scans: estimation of multi-vertex activation patterns

For each ROI from each subject, we separately estimated the multi-vertex activity pattern evoked by each stimulus and by each category in the task scans using two general linear models (GLM). One model used stimulus-specific β weights to estimate the multi-vertex activity pattern evoked by each individual stimulus, and the other used category-specific β weights to estimate the stimulus-evoked activity pattern associated with each category (e.g. the pattern outlined by the red square in Fig. 4A). Each GLM also included a separate target regressor for trials in which a stimulus was perturbed in size or position, and baseline and linear trend regressors for each scan. In order to determine the task-evoked magnitude for each stimulus category, a β weight matrix was separately computed using spatially non-normalized BOLD timeseries from the task scans. As with the localizer scans, all GLMs for the task scans were constructed using an assumed hemodynamic response.

### Task scans: Representational Similarity Analysis of stimulus-evoked patterns

We conducted a representational similarity analysis to verify that the estimated multi-vertex patterns for exemplars of a category were more similar to the patterns for exemplars from the same category vs. a different category. A representational similarity matrix (RSM) for each subject was computed by correlating across vertices the obtained β weights for each stimulus exemplar with the weights for each other stimulus exemplar. The individual RSMs were then averaged across all 16 subjects, with Fisher-Z transformations and reverse Fisher-Z transformations, to produce a group-averaged RSM (Fig. 6A). We conducted a second representational similarity analysis to determine if the multi-vertex patterns for each category showed the expected cross-category relationships, such as greater similarity between the patterns for the face and body categories than face and scene categories. For each individual an RSM was computed by correlating across vertices the obtained categorical β weights (e.g. the average scene-evoked multi-vertex pattern outlined by the red square in Fig. 4A) across categories, i.e. we correlated the multi-vertex pattern for scenes with the multi-vertex patterns for phase- or grid-scrambled, tools, chairs, mammals, bodies, and faces. A group-averaged RSM was then computed (Fig. 6B).

**Figure 6.**
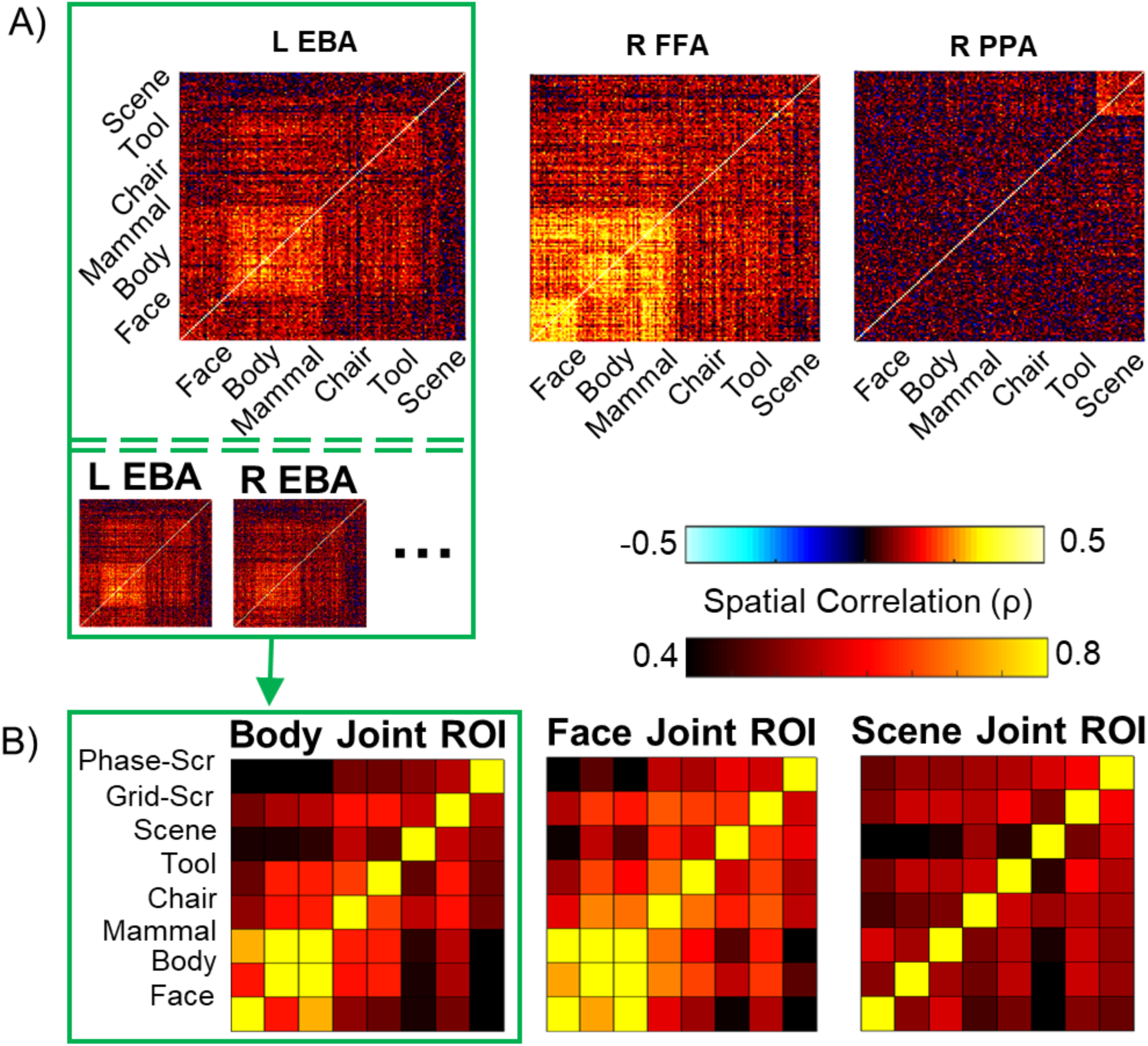
Representational similarity analysis (RSA) of individual stimuli and categories. **(A)** RSA based on the multi-vertex patterns for individual exemplars for each ‘whole-object’ category in 3 classical category-preferential areas. **(B)** RSA based on the multi-vertex pattern evoked for each category.

### Determining similarity of resting multi-vertex patterns and stimulus-evoked patterns

The next step in the analysis determined the similarity of stimulus-evoked multi-vertex patterns to the patterns observed on each frame of resting scans. For each participant’s individual joint-ROI and the associated constituent ROIs, we determined the degree to which the stimulus-evoked multi-vertex pattern for a category matched the multi-vertex pattern on each resting frame. The procedure is illustrated in Figure 7A for a single subject using real data. In the first step, as described above, the multi-vertex pattern evoked by a category in a region was determined (e.g. the ‘Scene’ activity pattern outlined by the red square in Fig. 7A, **‘Task BOLD’**). Then, the stimulus-evoked pattern for a category was correlated across vertices with the resting activity pattern on a single frame of the resting-state scans (Fig. 7A, **‘Resting-state BOLD’)**. A high positive correlation coefficient indicates that the resting multi-vertex pattern on a given frame was very similar to the pattern evoked by the category (e.g. the resting frame with a ‘Scene’-like resting activity pattern outlined by the magenta square in Fig. 7A). A near zero correlation coefficient indicates that the resting multi-vertex pattern on a given frame was not similar to the pattern evoked by the category (e.g. the resting frame with a not-‘Scene’-like resting activity pattern outlined by the green square in Fig. 7A). Finally, a high negative correlation coefficient indicates that the resting multi-vertex pattern on a given frame was very similar to the inverse of the pattern evoked by the category (e.g. the resting frame with a ‘Scene’-inverted resting activity pattern outlined by the cyan square in Fig. 7A). This procedure was repeated across all resting frames, resulting in a ‘stimulus-pattern-to-rest’ correlation timeseries (one correlation coefficient per resting frame) for each category in each ROI, as shown by the timeseries in Figure 7A.

**Figure 7.**
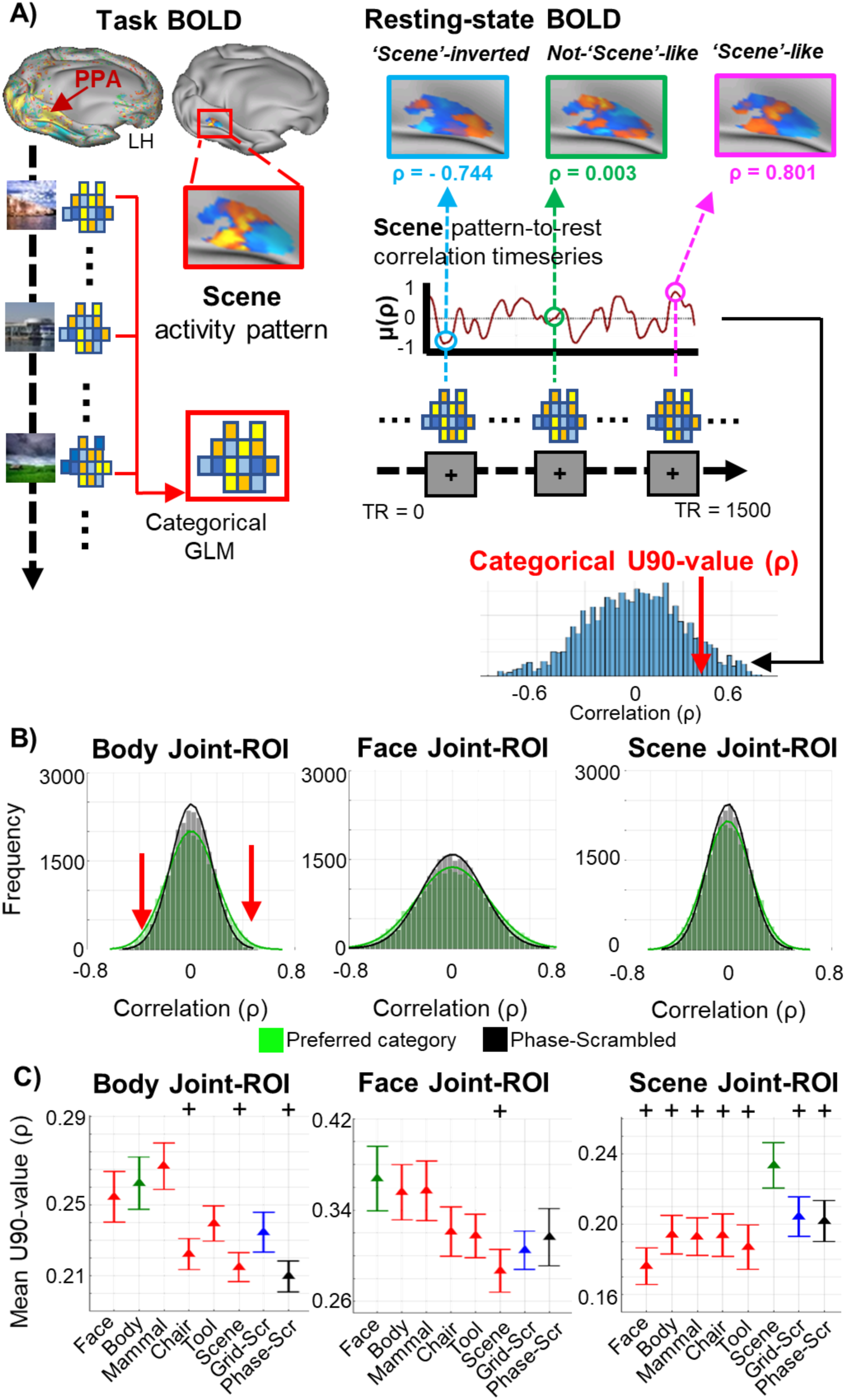
Stimulus-evoked-to-rest pattern similarity analysis in visual regions preferring specific categories. **(A)** Procedure for computing U90-values: determining the category-evoked multi-vertex pattern on task scans, correlating that pattern over vertices with the pattern on each resting frame, computing a U90 value from the resulting distribution of correlation coefficients. **(B)** Superimposed distributions of correlation coefficients for a joint-ROI’s preferred stimulus category, which was defined by the corresponding localizer contrast (light green; e.g. Body in the Body-preferred joint-ROI) and the phase-scrambled category (gray). **(C)** Group-averaged U90 values for the joint-ROI’s preferred category (green symbol), other whole-object categories (red symbols), grid-scrambled category (blue symbol), and phase-scrambled category (gray symbol). Black symbols indicate significant paired t-test between the joint-ROI’s preferred category and indicated category (++ = Bonferroni-Holm corrected p-val ≤ 0.005). Error bars indicate ±SEM.

From each timeseries, we constructed a corresponding distribution of correlation coefficients (Fig. 7A, distribution shown in blue). The upper 90% value of each distribution, hereafter termed the **U90-value**, was then determined (Fig. 7C). The U90 value computed for a category and ROI served as a measure of the relationship between resting activity patterns and the patterns evoked by a category mean. The U90-value was used as an alternative measure of variance since the U90-value refers to a correlation coefficient value, which indicates the degree of pattern similarity between the task-evoked and resting state activity pattern. Similar results were found using the variance of the distribution as a summary measure instead of the U90 value. For analyses that involved the Whole-Objects joint-ROI rather than joint-ROIs that preferred a particular object category such as faces, U90-values for the six whole-object categories (face, body, mammal, chair, tool, scene) were averaged together to form a whole-objects U90 value (Fig. 8B).

**Figure 8.**
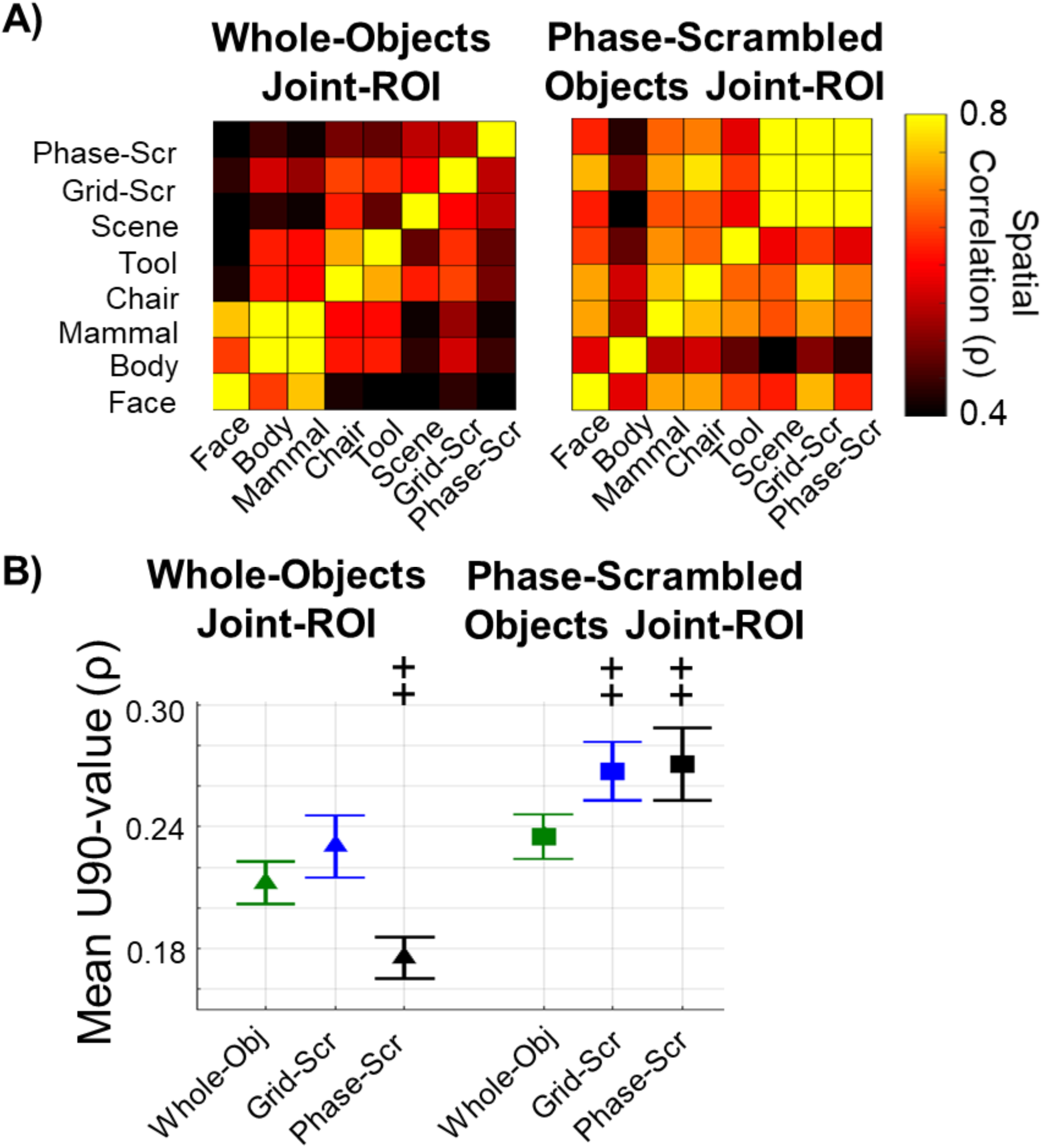
Stimulus-evoked-to-rest pattern similarity analysis in regions preferring phase-scrambled objects or whole-objects. **(A)** RSAs for Whole-Objects and Phase-Scrambled Objects joint-ROIs based on the multi-vertex pattern for each category. **(B)** Group-averaged U90 values. Black symbols indicate significant paired t-test between the whole-object category and scrambled category (++ = Bonferroni-Holm corrected p-val ≤ 0.005). Error bars indicate ±SEM.

Finally, to observe a potential relationship in a joint-ROI between the magnitude of the category-evoked response (Fig. 9A) and the strength of the relationship between category-evoked multi-vertex patterns and spontaneous activity patterns (U90 values), the correlation across category between task activation magnitudes and U90 values of stimulus-evoked-to-rest pattern similarity was measured for each joint-ROI (Fig. 9B).

**Figure 9.**
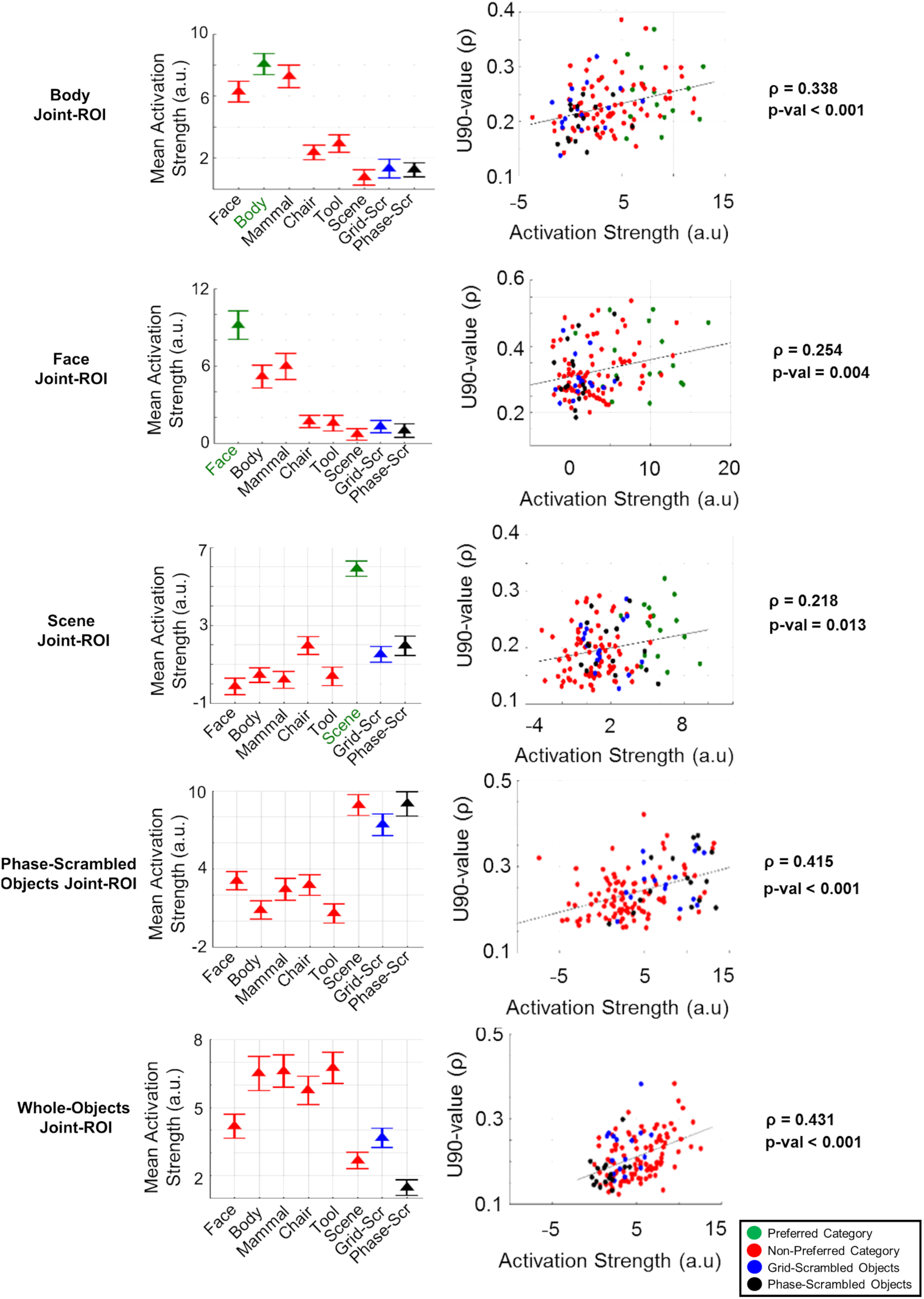
(A) Group mean activation strength from task scans for all stimulus categories in each joint-ROI. Error bars indicate ±SEM. **(B)** Correlation between activation strengths from task scans and U90 values across all categories and participants.

### Statistical analysis of U90 values

To statistically analyze the similarity between stimulus-evoked and resting multi-vertex patterns, U90 values were analyzed via repeated measures ANOVAs and post-hoc paired t-tests. For example, the statistical significance of an overall dependence of U90 values for a joint-ROI on the stimulus category was determined by conducting repeated-measures ANOVAs with Category-Type as factors. Paired t-tests were then conducted to test specific contrasts, with a Bonferroni-Holm correction for multiple comparisons.

### Pattern-based resting functional connectivity

The preceding analyses examined the similarity between stimulus-evoked patterns and the patterns measured on single frames of resting scans. The analyses described next determined the temporal correlation over resting frames of the amplitude of stimulus-evoked patterns in two different regions (pattern-based FC). For the pattern-based FC analysis, we used the constituent ROIs from the joint-ROI that preferred a category (face, body, or scene) relative to the object category (chairs + tools). Since only two face constituent ROIs were found and one of those ROIs largely overlapped with a Body constituent ROI in ventral temporal cortex, Face constituent ROIs were not included in the FC analysis. Therefore, pattern-based FC was computed over 14 ROIs: 5 from the Body joint-ROI and 9 from the Scene joint-ROI.

The correlation over vertices of a stimulus-evoked pattern with the pattern on a single resting frame results in a single correlation coefficient. Repeating this process for each successive resting frame results in a timeseries of correlation values, which we call a stimulus-pattern-to-rest correlation timeseries (see ***Determining similarity of resting multi-vertex patterns and stimulus-evoked patterns*** and Fig. 7A). Three pattern-based FC matrices were computed using stimulus-pattern-to-rest correlation timeseries. Figure 10A illustrates the procedure for computing the cells of an FC matrix using the stimulus-pattern-to-rest correlation timeseries from two scene regions (TOS, PPA) and two body regions (EBA, FBA). Figure 10B shows the resulting matrices. First, for each participant, stimulus-pattern-to-rest correlation timeseries for each of the 14 ROIs were generated using only the body-evoked pattern for each ROI (i.e. the multi-vertex activity pattern evoked by bodies in that ROI during the Task scans). The correlation between these body-pattern-to-rest timeseries for all pairings of the 14 ROIs was then computed (i.e. body-ROI-to-body-ROI, scene-ROI-to-scene-ROI, and body-ROI-to-scene-ROI pairings). For example, the leftmost graphs in Figure 10A show the correlation between TOS and PPA (top graph) and the correlation between EBA and FBA (bottom graph) using the body-pattern-to-rest correlation timeseries generated in each ROI from the body-evoked pattern. The correlation coefficients were then entered into the corresponding cells of the pattern-based FC matrix, “Using Body pattern-to-rest correlation timeseries only”, shown in Figure 10B.

**Figure 10.**
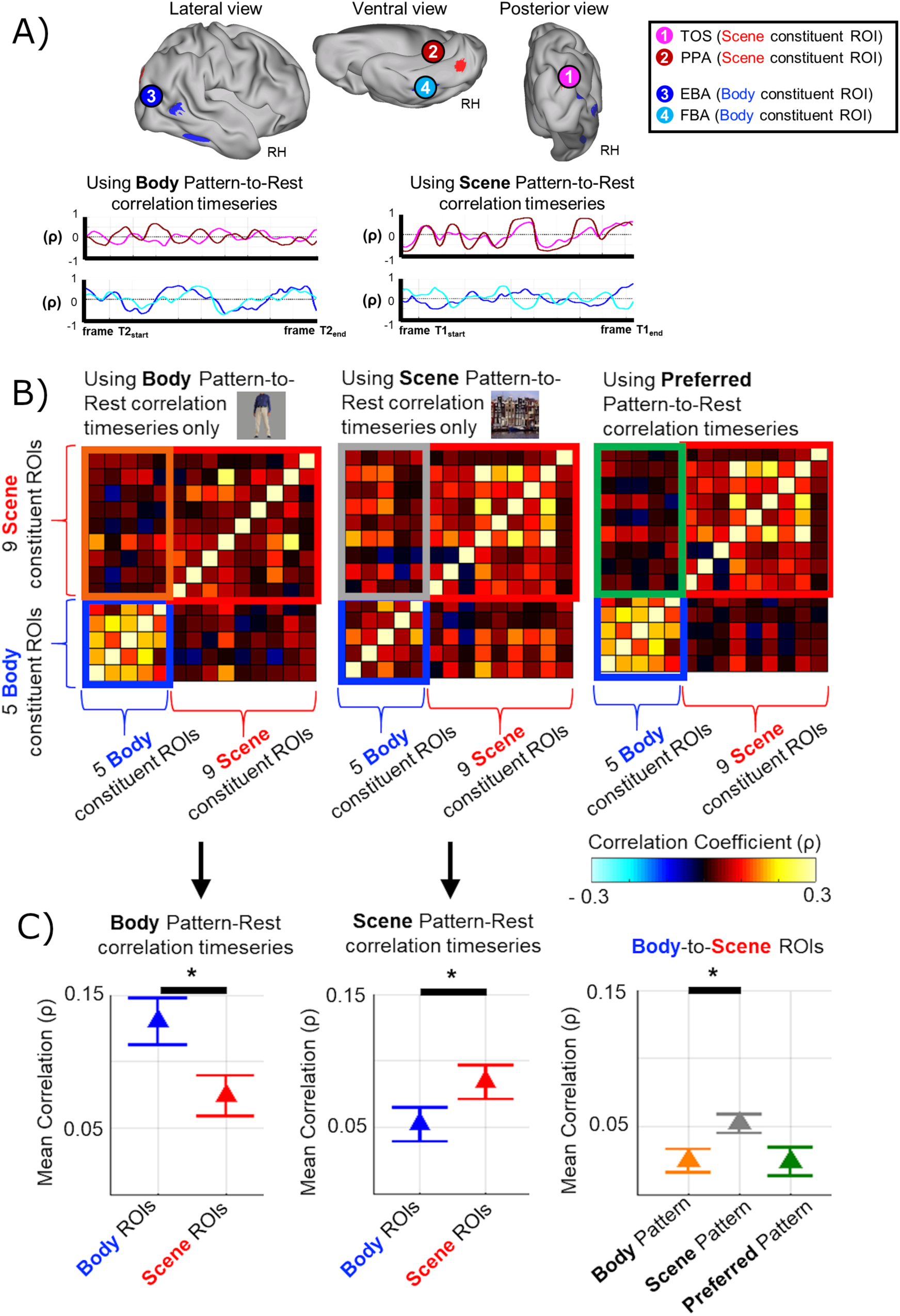
Pattern-based FC. **(A)** Stimulus-pattern-to-rest correlation timeseries computed using body-evoked and scene-evoked activity patterns in two scene-preferring and two body-preferring ROIs. **(B)** Pattern-based FC matrices computed using body-evoked, scene-evoked, or preferred category-evoked multi-vertex patterns (see text for details). **(C) Left, Middle**: Group-averaged pattern-based FC between body-preferring regions and between scene-preferring regions computed using body-evoked patterns (left) or scene-evoked patterns (middle). **Right**: Group-averaged pattern-based FC between body and scene regions computed using body-evoked, scene-evoked, or preferred-category-evoked patterns. Black symbols indicate significant group paired t-test comparing correlation (ρ) values (* = p-val ≤ 0.05). Error bars indicate ±SEM.

A similar procedure was used to generate a pattern-based FC matrix using only the scene-evoked pattern for each ROI. For example, the middle graphs in Figure 10A show the correlation between TOS and PPA, and between EBA and FBA using the stimulus-pattern-to-rest correlation timeseries generated in each ROI using the scene-evoked pattern. Finally, the correlation coefficients were entered into the corresponding cells of the pattern-based FC matrix, “Using Scene pattern-to-rest correlation timeseries only”, shown in Figure 10B.

To generate the third pattern-based FC matrix in Figure 10B (“using Preferred Stimulus pattern-to-rest correlation timeseries), the stimulus-pattern-to-rest correlation timeseries in a Body ROI was generated using the body-evoked pattern for that ROI and the stimulus-pattern-to-rest correlation timeseries in a Scene ROI was generated using the scene-evoked pattern for that ROI. Then the correlation between the correlation timeseries for all pairings of the 14 ROIs was computed and entered into the appropriate cells of the pattern-based FC matrix.

Finally, a vertex-averaged FC matrix was computed by first averaging the resting BOLD timeseries across all vertices of an ROI to generate a vertex-averaged timeseries, and then temporally correlating these averaged timeseries for all pairs of ROIs (Fig. 11). Vertex-averaged FC matrices, which correspond to the standard regional FC matrices found in the literature, eliminate any information carried by the multi-vertex pattern of BOLD activity within ROIs.

**Figure 11.**
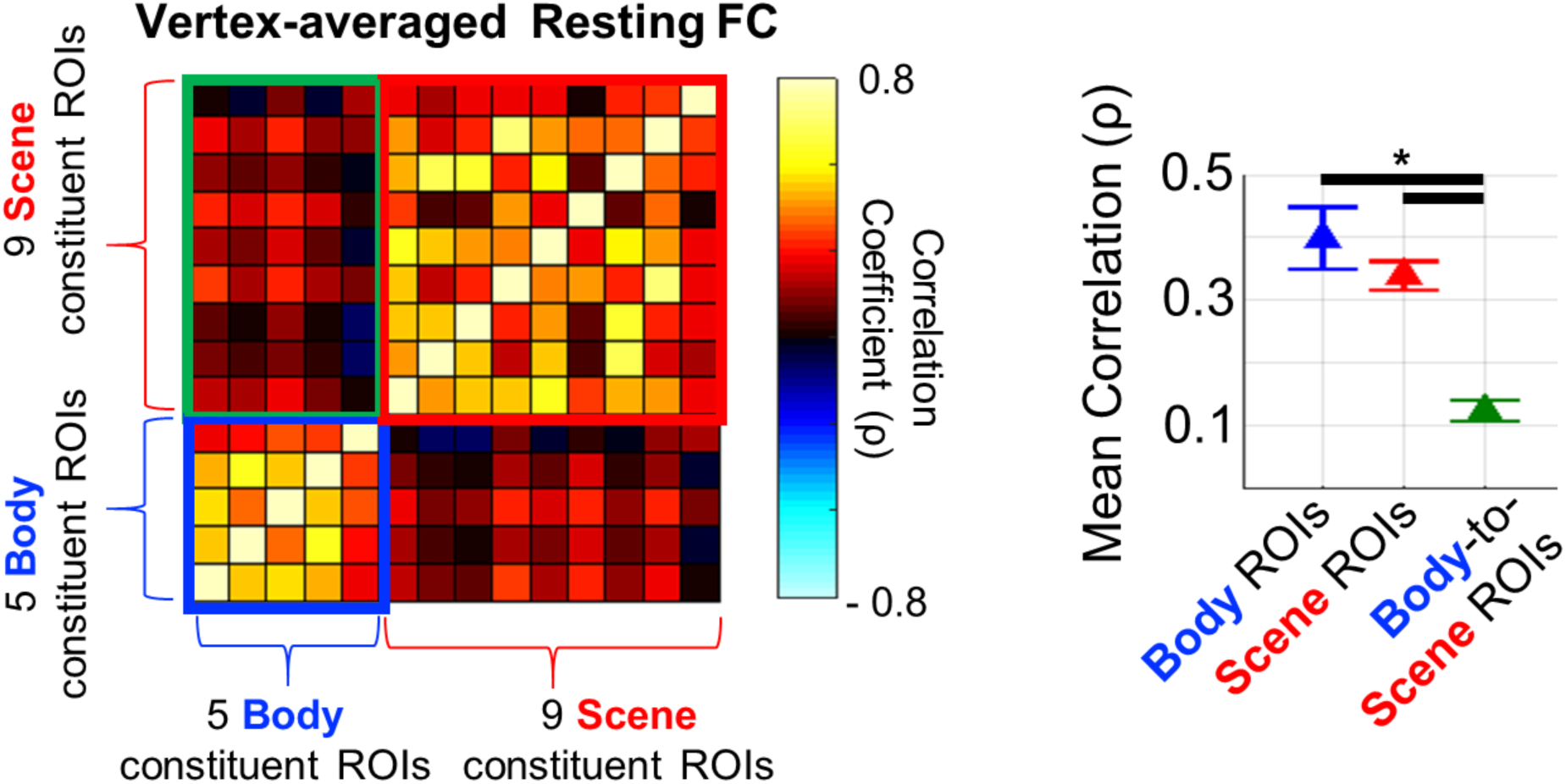
Vertex-averaged FC. A standard non-pattern-based FC matrix was computed by first averaging the vertex-wise timeseries of the BOLD signal across the vertices within each region, and then correlating the resulting vertex-averaged timeseries across all pairs of regions (**Left**). All cells involving FC between body regions, between scene regions, and between body and scene regions are averaged and plotted **(Right)**.

Pattern-based FC values were analyzed via repeated measures ANOVAs and paired t-tests. For example, we statistically evaluated whether the magnitude of pattern-based FC depended on both the category of the stimulus-evoked multi-vertex activity pattern and the preferred category of the ROIs by conducting a repeated-measures ANOVA with the Category-evoked pattern (body, scene) and ROI-Type (body, scene) as factors. Paired t-tests were conducted to test differences between specific evoked patterns/ROI combinations. For example, pattern-based FC values for body ROIs vs. scene ROIs were compared for correlation timeseries generated using body-evoked multi-vertex patterns.

## Results

The first goal of the experiment was to compare multi-vertex activity patterns measured in the resting state with fMRI to stimulus-evoked patterns for different stimulus categories, including ecological stimuli (e.g. photographs of faces, tools, and scenes) and stimuli that emphasized low-level features (e.g. phase-scrambled or grid-scrambled images of those stimuli). This comparison was conducted in regions of higher-order visual cortex that activated more strongly to specific stimulus categories (e.g. bodies) relative to other categories (e.g. chairs and tools). In addition, resting and stimulus-evoked patterns for phase-scrambled, grid-scrambled, and whole-objects were compared in regions of visual cortex that responded more strongly to phase-scrambled objects than to whole-objects or showed the reverse relationship. To measure spontaneous activity, we ran a set of resting-state scans in which human observers fixated a central point on a blank screen. This activity was measured first to prevent possible learning effects from the other conditions (Fig. 3).

### Localization of regions with visual category preferences

To identify category-specific visual regions, we ran a set of localizer scans in which multiple stimuli belonging to one of five stimulus categories (faces, bodies, scenes, man-made objects (chairs and tools), and phase-scrambled versions of these stimuli) were presented in a blocked design (Figs. 2 and 3). We used standard contrasts (as in (Bracci & Op de Beeck 2016)) to identify category-preferring regions. For instance, activity evoked by body stimuli was subtracted from activity evoked by man-made objects (tools and chairs) to localize body-preferring regions such as EBA (see Figs. 4A **and** 4B, and Table 1 for all category-preferring regions). Separate contrasts identified regions more active for whole-objects (face+scene+bodies+(tools+chairs)) than for low-level visual features (phase-scrambled objects). Phase-scrambled objects activated more strongly in regions of early visual cortex (V1-V3 based on the maps of (Wang et al 2015)), while whole objects activated more strongly in lateral and ventral occipital cortex, including some category-preferring regions (Figs. 5A and 5B, Table 1).

### Representational similarity analysis of task-evoked patterns

During task scans (Fig. 3), we randomly presented individual stimuli belonging to each category to extract the stimulus-evoked multi-vertex pattern in a particular ROI for each stimulus and corresponding category. Two general linear models (GLMs) were conducted to estimate the stimulus-evoked patterns. One model used stimulus-specific β weights to estimate the multi-vertex activity pattern evoked by each individual stimulus, and the other used category-specific β weights to estimate the stimulus-evoked multi-vertex activity pattern associated with each category.

To show that our stimuli and procedure generated multi-vertex patterns that were consistent with the literature, we conducted a representational similarity analysis (RSA). The representational similarity analysis was conducted using the task scans, which were completely independent of the localizer scans used to determine category-preferring ROIs. Figure 6A shows the similarity of multi-vertex patterns evoked by individual stimuli within several classical category-preferring ROIs. In left EBA the highest representational similarity was found between human bodies, and the next highest between pictures of mammals, which included their bodies. In the right fusiform face area (FFA (Kanwisher et al 1997)), faces and other animate stimuli (bodies, mammals) generated more similar patterns than stimuli from the inanimate categories chair, tool, and scene, with the most consistent representational similarity found between face exemplars (Grill-Spector & Weiner 2014, Kriegeskorte et al 2008). In the scene-preferring region right parahippocampal place area (PPA (Epstein & Kanwisher 1998)), the activity patterns evoked by different scenes were well correlated, with low correlations between and within all other categories.

We conducted a second representational similarity analysis using the pattern evoked by a stimulus category, as estimated by a category regressor in a separate GLM, rather than using the patterns evoked by individual stimuli. In addition, we grouped each set of category-preferential ROIs for an individual into a single joint-ROI instead of conducting the analysis separately within each localizer-defined ROI. For instance, the body joint-ROI included left and right EBA, left and right fusiform body area (FBA), and so forth, and the scene joint-ROI included constituent regions such as PPA, the transverse occipital sulcus (TOS), and retrosplenial cortex (RSC) (see Table 1 for a complete listing). Therefore, a joint-ROI included all the vertices from visual regions that preferred a certain visual category. This procedure simplified the analysis of the similarity between stimulus-evoked and resting multi-vertex patterns (see ***Methods***, ***Defining ROIs from localizer activation contrasts*** for the full rationale behind the use of joint-ROIs).

The results of this second RSA procedure (Fig. 6B) were also consistent with the literature. Both in body and face joint-ROIs, the highest representational similarity was found between animate categories (face, bodies, mammals) as compared to other categories (Grill-Spector & Weiner 2014, Kriegeskorte et al 2008). For instance, the similarity of body- and face-evoked multi-vertex activity patterns in the face joint-ROI was ρ=0.73, while the similarity of face- and scene-evoked patterns in the face joint-ROI was ρ=0.41. Table 2 indicates the representational similarity between the task-evoked multi-vertex patterns corresponding to the face, body, and scene categories within each joint-ROI.

**Table 2.** Correlation over vertices between face-evoked, body-evoked, and scene-evoked multi-vertex patterns in three joint-ROIs.

| | Similarity ($\rho$) between<br>Face-evoked activity pattern<br>and<br>Body-evoked activity pattern | Similarity ($\rho$) between<br>Face-evoked activity pattern<br>and<br>Scene-evoked activity pattern | Similarity ($\rho$) between<br>Body-evoked activity pattern<br>and<br>Scene-evoked activity pattern |
| --- | --- | --- | --- |
| Face Joint-ROI | 0.729 | 0.415 | 0.550 |
| Body Joint-ROI | 0.618 | 0.421 | 0.423 |
| Scene Joint-ROI | 0.506 | 0.356 | 0.379 |

### Stimulus-evoked-to-rest pattern similarity analysis in category-preferring regions

We next tested the first prediction of the representation hypothesis, namely that multi-vertex patterns of spontaneous activity in category-preferring regions should be more related to the stimulus-evoked multi-vertex pattern for preferred than non-preferred categories. For each joint-ROI and for each category, the category-evoked pattern for the joint-ROI was correlated over the vertices of the joint-ROI with the resting pattern in the joint-ROI on each resting frame to determine a resting timeseries of correlation coefficients (termed a stimulus-pattern-to-rest correlation timeseries) and a corresponding frequency distribution of coefficient values. The upper 90% value (U90 value) of the distribution was used as a summary measure of the relationship between the stimulus-evoked and resting multi-vertex activity patterns (see **Methods**, ***Determining similarity of resting multi-vertex patterns and stimulus-evoked patterns*** for the rationale behind using U90 values as opposed to measures of variance).

Figure 7A illustrates this procedure in a single subject using a single ROI that prefers scenes (PPA) and the category-evoked pattern for scenes, which is a multi-vertex set of normalized activation values (Fig. 7A, left). This evoked pattern is correlated over the vertices of PPA (ρ) with the spontaneous patterns of activity on each frame of the resting state scans. A timeseries of ρ-values is generated (Fig. 7A, middle), as well as a corresponding frequency distribution of ρ-values (Fig. 7A, the histogram in blue). A U90 value for the ROI and category is then determined from the distribution. The insets in the middle panel show resting frames in which the spontaneous activity (real data) was not correlated (ρ=0.003, outlined in green), positively correlated (ρ=0.81, outlined in magenta), or negatively correlated (ρ=-0.74, outlined in cyan) with the scene-evoked multi-vertex pattern. The same procedure was used to generate a U90 value for each of eight categories (faces, bodies, mammals, chairs, tools, scenes, grid-scrambled, phase-scrambled) in each joint-ROI (body, face, scene).

Figure 7B shows the distributions of correlation coefficients across all subjects for each preferred category within its corresponding joint ROI (green hue). For example, the leftmost graph shows the distribution using the body-evoked multi-vertex pattern within the body joint-ROI. A second distribution, generated using the pattern evoked by phase-scrambled objects within the same body joint-ROI, has been superimposed (black hue). Theoretically, the two distributions might differ in the mean, variance, skewness, or some other parameter. For each joint-ROI, the distribution of correlation coefficients for both the preferred stimulus category and the phase-scrambled category were symmetric and centered on zero. However, the spread of the distribution was higher for the preferred category-evoked multi-vertex pattern, meaning that larger correlation coefficients, both positive and negative, were observed for the preferred than phase-scrambled category (red arrows in Fig. 7B). Therefore, a larger U90 value indicates the presence of larger positive matches and larger negative matches between the resting multi-vertex pattern and the category-evoked pattern. Similar findings were obtained for face (middle panel) and scene multi-vertex patterns (rightmost panel) in the corresponding joint-ROIs, as compared to phase-scrambled patterns.

The categorical specificity of spontaneous multi-vertex patterns in each joint-ROI was tested by comparing U90 values for different categories. Figure 7C shows mean U90 values for a joint-ROI’s preferred category, defined from its localizer contrast (green symbol; e.g. body in the Body joint-ROI), “non-preferred” categories (red symbols), grid-scrambled category (blue symbol) and phase-scrambled category (black symbol) averaged across subjects. We first conducted an overall repeated measures analysis of variance ANOVA on U90 values with joint-ROI (body, face, and scene) and Category (8 levels) as factors. The main effects of joint-ROI (F(2, 30)=50.4, p<.0001) and Category (F(7, 105)=3.46, p=.002), and the interaction of joint-ROI by Category (F(14, 210)=4.37, p<.0001) were all significant. Separate repeated measures ANOVAs for each joint-ROI with Category (8 levels) as a factor indicated a highly significant main effect of Category in each joint-ROI (Body: F(7,105)=7.25, p<.0001; Face: F(7,105)=3.25, p=.004; Scene: F(7,105)=3.93, p=.0008). Therefore, for each joint-ROI, the spread of stimulus-pattern-to-rest similarity values significantly depended on the category of the stimulus-evoked pattern.

The significant main effect of Category for each joint-ROI indicated that resting multi-vertex patterns in each joint-ROI consistently showed larger U90 values for some category-evoked patterns than for other patterns. The significant interaction of joint-ROI by Category indicated that these modulations of U90 values across category reliably differed across joint-ROIs. These results are consistent with representation hypothesis.

To compare the U90 value for the joint-ROI’s preferred category vs. each other category, we conducted paired t-tests with a Holm-Bonferroni correction for multiple comparisons. Significant, multiple-corrected comparisons are indicated in Figure 7C by plus signs. In the Body joint-ROI, the U90 value for bodies was significantly larger than for chairs, scenes, and phase-scrambled stimuli. In the Face joint-ROI, the U90 value for faces was significantly larger than for scenes. Conversely, in the Scene joint-ROI, the U90 value for scenes was significantly larger than for all other categories.

Therefore, the ‘animate’ Body and Face joint-ROIs and the ‘inanimate’ Scene joint-ROI showed a significant double dissociation involving the corresponding categories, with U90 values in the Face and Body joint-ROIs significantly greater for the face and body categories, respectively, than for scene categories, and the U90 value in the inanimate Scene joint-ROI significantly greater for scenes than for either faces or bodies. The U90 values for face and body categories within each joint ROI were similar, reflecting the fact that both are animate categories and have greater cross-category representational similarity with each other than with scenes (Table 2). These results provide some support for the first prediction of the representation hypothesis, namely that spontaneous multi-vertex patterns in category-preferring regions are more related to the patterns for some categories than for others. However, ‘more related’ means a greater spread of extreme similarity values, both positive and negative, rather than a shift in the mean to more positive similarity values.

### Stimulus-evoked-to-rest pattern similarity analysis in regions preferring whole vs. phase-scrambled objects

The above results showed that the multi-vertex pattern of spontaneous activity in regions of high-level visual cortex that respond preferentially to ecological visual categories was more related to the pattern evoked by one category than another (e.g. bodies vs. scenes). We next asked whether a similar result would be found in regions that show stimulus preferences for low-level features as compared to more ecological categories such as face or body. This result would support a general conclusion that the stimulus preferences of a region largely drive the multi-vertex pattern of spontaneous activity. We used the localizer scans to identify ROIs in which stimulus-evoked responses were stronger or weaker for phase-scrambled objects than for the union of the whole-object categories (face, body, mammal, chair, tool, and scene). The resulting ‘Phase-scrambled objects’ joint-ROI comprised medial posterior visual regions in early visual cortex (V1-V3 according to the Wang template (Wang et al 2015) while the ‘Whole-objects’ joint-ROI comprised regions in lateral and ventral visual cortex (Fig. 5B).

A representational similarity analysis in the Phase-scrambled objects joint-ROI showed high similarity between phase-scrambled, grid-scrambled, and scene stimuli, while the Whole-objects joint-ROI showed low similarity between those categories (Fig 8A). Figure 8B shows the results of a stimulus-evoked-to-rest pattern similarity analysis based on U90 values in each joint-ROI, which support the general conclusion that stimulus-evoked-to-rest pattern similarities are not necessarily stronger for more ecological stimuli. Instead, stimulus-evoked-to-rest correspondences reflect a regions’ stimulus preferences, which are different in high- and low-level visual cortical regions (Fig. 8B). An ANOVA with ROI-type (Whole-objects, Phase-scrambled objects) and Stimulus-type (whole-objects, grid-scrambled, phase-scrambled objects) as factors indicated that the critical interaction of ROI-type by Category (F(2,30)=14.2, p<.0001) was significant. A significant interaction was also found for a 2 x 2 sub-ANOVA restricted to the categories whole objects and phase-scrambled objects (F(1,15)=23.5, p<.0001). These results are again consistent with representation hypothesis.

Within each joint-ROI, we compared the U90 value for the preferred category vs. the two “non-preferred” categories using paired t-tests with a Holm-Bonferroni correction for the four comparisons over the two joint-ROIs. In the Phase-scrambled joint-ROI, U90 values were significantly higher for both scrambled stimulus categories than for the whole-objects category, and in the Whole-objects joint-ROI, the U90 value for the whole-object category was significantly greater than for the phase-scrambled object category. The grid-scrambled pattern, which contains both high-level and low-level features (e.g. a high density of contour terminators), showed U90 values both in early visual and higher-order visual cortex that were not distinguishable from the regions’ preferred stimulus category.

These results demonstrate a second double dissociation relating the dependence of U90 values on both the category-evoked multi-vertex pattern and the joint-ROI in which the similarity of the evoked pattern to spontaneous patterns was evaluated. They are consistent with the interpretation that spontaneous activity patterns in visual cortex are strongly affected by the stimulus preferences of the region, irrespective of whether those preferences favor more or less ecological categories.

### U90 values correlate with activation strength

The representation hypothesis maintains that task-evoked patterns entrain spontaneous activity patterns during development and through experience. Therefore, one might expect a positive relationship between the magnitude of the category-evoked response and the strength of the relationship between category-evoked multi-vertex patterns and spontaneous activity patterns (i.e. the U90 value). Figure 9A shows the mean activation strengths for different categories during the Task scans. The magnitude of the stimulus-evoked response in a joint-ROI was generally strongest for the preferred category. Since joint-ROIs were defined from localizer scans that were independent of the task scans, this result indicates the stability of the ROI assignments.

Figure 9B shows the correlation across category between task activation magnitude and U90 values of stimulus-evoked-to-rest pattern similarity for each joint-ROI. There was a positive and significant correlation between task activation values and U90 values in each joint-ROI. The greater the activation strength of a category, the greater the U90 value, a relationship that held for joint-ROIs irrespective of their stimulus preferences. This relationship also was significant when the correlation coefficient between activation strength and U90 value was computed separately for each participant, and a group 1-sample t-test was conducted. Correlation coefficients were highly significant in all joint-ROIs (Body, p=.0004, Face, p=.0047, Scene, p=.0014; Whole-object, p=.0002; Phase-scrambled, p<.0001).

### Pattern-based functional connectivity at rest

FC analyses typically evaluate the correlation between the timeseries of activity for single voxels or between voxel-averaged timeseries. However, recent studies have also measured the inter-regional temporal correlation of multi-vertex patterns during tasks (Anzellotti & Coutanche 2018, Chen et al 2018, Coutanche & Thompson-Schill 2013). We used a similar approach to determine whether the signed magnitude of the resting multi-vertex pattern for a category in each constituent ROI of a joint-ROI, as determined on a resting frame by correlating over vertices the category-evoked and resting activity pattern, fluctuated synchronously or independently over frames between pairs of constituent ROIs. For instance, we determined whether the amplitude of the multi-vertex patterns for scenes in regions such as PPA, TOS, and RSC, which were previously combined to form the Scene joint-ROI (Table 1), fluctuated synchronously at rest. Synchronous fluctuations (i.e. temporally correlated fluctuations) would indicate temporal variations of the amplitude of an inter-regional brain state specific for a particular category. We conducted this analysis across the constituent ROIs of the Body and Scene joint-ROIs. The Face joint-ROI was not included in these analyses, since that ROI only included two regions and one of them overlapped with a constituent body ROI. In contrast, the Scene and Body joint-ROIs contained multiple ROIs that were all disjoint.

For each body- and scene-preferring constituent ROI, separate body and scene “stimulus-pattern-to-rest” correlation timeseries were computed based, respectively, on the correlation over vertices of the multi-vertex pattern on each resting frame with the body-evoked pattern and the scene-evoked pattern. This procedure is illustrated in Figure 10A using data from segments of resting scans in one subject. Stimulus-pattern-to-rest-correlation timeseries are shown for two scene-preferring regions (right PPA and right TOS) and two body-preferring regions (right EBA and right FBA). Each stimulus-pattern-to-rest-correlation timeseries shows the similarity values over time of resting multi-vertex patterns to a category-evoked pattern in a single ROI. For example, the left lower dark blue timeseries in Figure 10A shows the similarity of the resting pattern on each frame to the body-evoked pattern in the body constituent region EBA. The left lower graph shows that the body-pattern-to-rest correlation timeseries for body-preferring regions (right EBA and right FBA), which were computed using body-evoked multi-vertex patterns, are positively correlated. In contrast, the left upper graph shows that the body-pattern-to-rest correlation timeseries for scene-preferring regions (right PPA and right TOS), which again were computed using body-evoked multi-vertex patterns, are uncorrelated. Conversely, when stimulus-pattern-to-rest correlation timeseries were computed using the scene-evoked multi-vertex pattern, the opposite results are found. The scene-pattern-to-rest correlation timeseries for scene-preferring regions show positively correlated fluctuations (right upper graph), while the scene-pattern-to-rest correlation timeseries for body-preferring regions are weakly correlated (right lower graph).

The data from all resting scans of all subjects were analyzed and the results are summarized in Figures 10B and 10C. Figure 10B shows three resting ‘pattern-based FC’ matrices. A pattern-based ‘body’ FC matrix (leftmost matrix) was constructed by computing all pairwise inter-regional correlations between the body-pattern-to-rest correlation timeseries, which were computed using body-evoked multi-vertex patterns. Similarly, a pattern-based ‘scene’ FC matrix (middle matrix) was computed using scene-pattern-to-rest correlation timeseries (i.e. timeseries computed using scene-evoked patterns). Qualitatively, body-preferring ROIs showed stronger positively correlated spontaneous fluctuations for body-evoked than scene-evoked patterns, and scene-preferring ROIs showed stronger positively correlated spontaneous fluctuations for scene-evoked than body-evoked patterns.

The graphs in Figure 10C summarize the pattern-based Body and Scene FC matrices by averaging the inter-regional correlations for body-preferring regions (the lower left block of each matrix in Fig. 10B outlined in blue) and scene-preferring regions (the upper right block of each matrix outlined in red). The left and middle graphs show respectively the results when stimulus-pattern-to-rest timeseries were computed using body-evoked and scene-evoked multi-vertex patterns. A 2-factor ANOVA on the mean pairwise pattern-based FC values with ROI-type (body, scene) and Category-evoked-pattern (body, scene) as factors yielded a main effect of ROI-type (F(1,15)=5.41, p=.035), reflecting the larger FC values in body-preferring regions and, critically, a significant interaction of ROI-type by Category-evoked pattern (F(1,15)=8.96, p=.009). This effect was further supported by paired t-tests of specific contrasts. When stimulus-pattern-to-rest timeseries were computed using a body-evoked multi-vertex pattern, inter-regional correlations were significantly higher in body-than scene-preferring ROIs. Conversely, when stimulus-pattern-to-rest timeseries were computed using a scene-evoked multi-vertex pattern, inter-regional correlations were significantly higher in scene-than body-preferring ROIs.

Therefore, spontaneous fluctuations of multi-vertex patterns of activity were more strongly correlated for multi-vertex patterns corresponding to the regions’ preferred category. This result suggests that resting pattern-based FC is modulated by the putative representational content of spontaneous activity. In addition, a paired t-test indicated that in the pattern-based Body FC matrix, the average FC was less in the scene-body block than in the body-body block of the matrix (p<.001; Fig. 10B, leftmost matrix, orange vs blue outlined blocks). Similarly, in the pattern-based Scene FC matrix, the average FC was less in the scene-body block than in the scene-scene block of the matrix (p=.035; Fig. 10B, middle matrix, gray vs red outlined blocks). Therefore, pattern-based FC was greater between regions preferring the same category than between regions preferring different categories.

A related question was whether putative body and scene representations fluctuated independently at rest. The rightmost matrix in Figure 10B shows a Preferred-Category pattern-based FC matrix in which body-pattern-to-rest correlation timeseries were computed in body-preferring regions (i.e. timeseries were computed using body-evoked multi-vertex patterns) and scene-pattern-to-rest correlation timeseries were computed in scene-preferring regions (i.e. timeseries were computed using scene-evoked multi-vertex patterns). Accordingly, the lower left and upper right blocks of the Preferred-Category matrix match, respectively, the lower left block of the Body pattern-based FC matrix and the upper right block of the Scene pattern-based FC matrix.

The ‘scene-body’ block of the Preferred Category matrix outlined in green is of primary interest. The correlation between scene and body regions was uniformly low under conditions in which the inter-regional correlation involved timeseries from scene and body regions that respectively indicated the fluctuations of scene- and body-evoked multi-vertex patterns (see rightmost graph, Fig. 10C, for average correlation values for scene-body blocks. Therefore, periods in which a body-evoked pattern was maximally present in body-preferring ROIs were largely independent of periods in which a scene-evoked pattern was maximally present in scene-preferring ROIs. Paired t-tests indicated that correlations in scene-body blocks from the Preferred-Category matrix were significantly lower than the correlations from scene-scene (p=.009) and body-body region blocks (p < .0001).

Finally, a standard FC matrix (Fig. 11, left panel) was constructed by computing vertex-averaged resting timeseries for each region, followed by pairwise correlation of the regional timeseries. As in previous work (Hutchison et al 2014, Konkle & Caramazza 2017, Wang et al 2016, Zhu et al), vertex-averaged FC was category specific, with stronger FC between body-preferring regions and between scene-preferring regions, than between body- and scene-preferring regions. Pattern-based FC matrices were moderately-to-strongly correlated with the vertex-averaged FC matrix. The largest correlation was with the preferred-category matrix rather than the matrices generated using a single category (body-evoked pattern, r=0.57; scene-evoked pattern, r=0.48; preferred-category, r=0.65).

### Category selectivity of U90 values in constituent ROIs

Figure 12 shows the category selectivity of U90 values for the individual constituent ROIs within a joint-ROI. For each joint-ROI, we conducted a two-factor ANOVA with Category and Constituent-ROI as factors and U90 value as the dependent measure. A main effect of Category with no interaction between Category and Constituent-ROI was observed for both the body joint-ROI (F(7,63)=2.45, p=.028) and the scene joint-ROI (F(7,63)=2.41, p=.03), indicating a consistent profile of U90 values over categories across the constituent ROIs of each joint-ROI. Nevertheless, variability in the category profiles over the constituent ROIs is evident. Since there were many fewer vertices in each constituent ROI than in the associated joint-ROI, some variability in category selectivity over constituent ROIs is expected due to noise. Additionally, however, we argue in the discussion that the category selectivity of U90 values for a joint-ROI is aided by the category selectivity of the pattern-based FC between its constituent ROIs.

**Figure 12.**
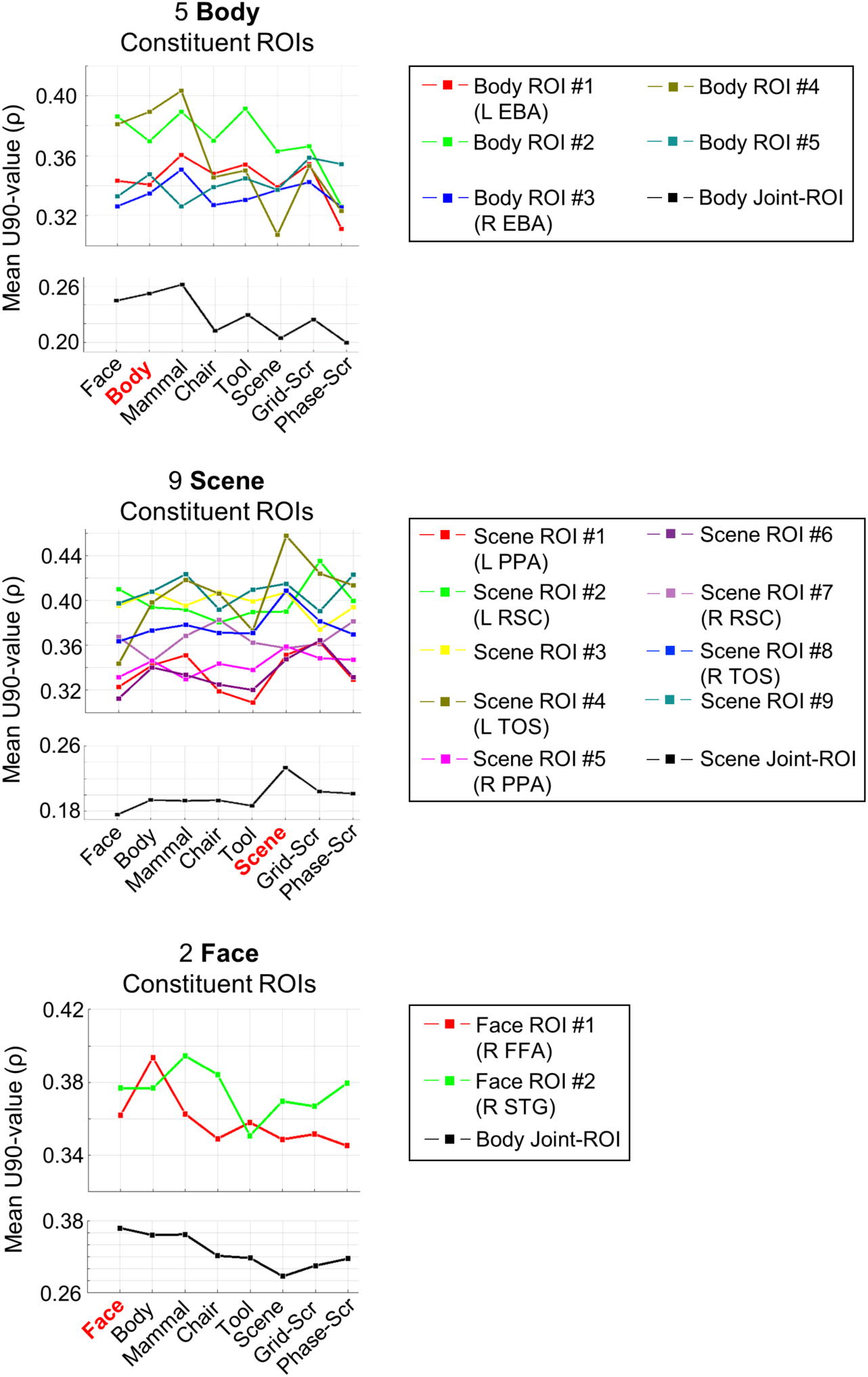
Graphs of the profile of group-averaged U90 values across stimulus categories for each constituent body region, scene region, and face region.

## Discussion

The goal of the experiment was to test representational theories of spontaneous activity by determining whether in regions of human visual cortex there is a link between multi-vertex patterns of spontaneous activity, measured in the resting-state, and the multi-vertex patterns evoked by ecological visual stimuli such as bodies or stimuli that emphasized low-level features such as phase-scrambled bodies.

We obtained two main results. First, resting multi-vertex activity patterns in regions of visual cortex were more closely related to the patterns evoked by the regions’ preferred stimulus categories. This relationship did not reflect a greater average similarity of resting patterns to the patterns for preferred categories, but instead a greater spread of resting similarity values for preferred categories. As resting multi-vertex patterns in a region fluctuated over time, we observed both larger positive and larger negative similarity values for the patterns evoked by stimulus categories preferred by the region. This result was demonstrated statistically by two significant double dissociations. Body- and face-preferring regions showed larger U90 values, indexing the spread of similarity values, for faces and bodies than for scenes, while scene-preferring regions showed larger U90 values for scenes than for faces and bodies (Fig. 7). Regions preferring whole objects vs. phase-scrambled objects showed a similar double dissociation (Fig. 8). This result was further strengthened by the positive correlation between U90 values and stimulus specific activation magnitudes. The more strongly a stimulus activated a region, the higher the spread of similarity values over resting frames between the stimulus-evoked multi-vertex pattern and the spontaneous pattern (Fig.9). The latter result is consistent with the notion that task-evoked patterns entrain spontaneous activity patterns during development and through experience (Fig. 1).

The second main result was that multi-vertex activity patterns evoked by a category fluctuated more synchronously at rest between cortical regions preferring that category. For example, in the resting-state the pattern evoked by bodies was more positively correlated between body-preferring ROIs than between scene-preferring ROIs. The pattern evoked by scenes showed the opposite result (Fig. 10). Finally, the multi-vertex patterns evoked by scenes and bodies fluctuated largely independently at rest within the preferred regions for those categories.

Therefore, the current results show that multi-vertex patterns of spontaneous activity within regions of human cortex, and fluctuations of those patterns between regions, code for stimulus and category specific information. In this respect, our work is more related to seminal work conducted in cats (Kenet et al 2003) and monkeys (Fukushima et al 2012, Omer et al 2018) than to previous work on category-selective visual regions in humans, which has focused on measurements of voxelwise functional connectivity rather than on measurements of the resting multi-vertex pattern of activity at a timepoint (Hutchison et al 2014, Stevens et al 2017, Strappini et al 2018, Turk-Browne et al 2010, Wilf et al 2017, Zhang et al 2009, Zhu et al). The most closely related previous work in humans was reported by Chen and colleagues, who compared voxelwise functional connectivity measurements to evoked multi-vertex patterns (Chen et al 2017).

### Spontaneous activity patterns for objects and features in visual cortex

Spontaneous activity patterns were not more similar on average to preferred stimulus evoked-patterns, but showed the greatest variation with respect to those patterns (Fig. 7). Interestingly, animal studies thus far have found the same result with some caveats. For instance, Kenet et al. (Kenet et al 2003) recorded voltage sensitive dye imaging in anesthetized cat visual cortex and found a significant correlation (r=0.6) between spontaneous activity patterns and orientation selective stimulus evoked patterns. Positive and negative values of the correlation distribution were higher (as in our experiment), rather than the mean, as compared to a control distribution obtained by flipping the orientation selective map. This finding was replicated recently in anesthetized monkey visual cortex (Omer et al 2018). In auditory monkey cortex, spontaneous spatial covariations of gamma activity recorded with cortical grids from auditory cortex resemble tonotopic maps derived with auditory stimuli. Also in this case, the correlation involves both positive and negative high correlation values as compared to a control distribution obtained by shuffling the tonotopic map (Fukushima et al 2012).

In our experiment, the control distribution was not spatially shuffled because this control might not preserve the local structure of the vascular architecture that is the anatomical basis of the measured BOLD signal. Therefore, we instead compared stimulus-evoked-to-rest correlation distributions for two different stimuli (e.g. bodies vs. scenes). Although the animal and human experiments differed in many ways, in all experiments the reported match between spontaneous and task-evoked activity was not a shift in the mean, but rather a higher frequency of more extreme matches/mismatches of the spatial patterns (e.g. Compare our Fig. 7B to (Omer et al 2018) Figs.1-2).

An issue for future work is whether spontaneous activity patterns are more related to the average pattern evoked by a category or the patterns evoked by individual exemplars from the category. In rat hippocampus, spontaneous replay in anesthetized and awake animals of sequences of activity during navigation, a phenomenon qualitatively similar to what reported here (e.g. (Karlsson & Frank 2009); see also (Buzsaki & Draguhn 2004)), seem to reflect individual experiences rather than averages. Hippocampal replay sequences have been mainly conceptualized as reflecting a mechanism for consolidating information in long-term memory.

One interpretation of the current results is that spontaneous activity serves as a prior for task processing such as object recognition. An important rationale for postulating a representational function of resting activity is that limits on the information processing capacity of the brain may be mitigated by the incorporation of useful prior information. Appropriate priors will generally depend on context and therefore will change dynamically. The perceptual priors appropriate to walking alone through a forest vs. eating a family meal at the dinner table are quite different. These putative dynamic changes are thought to reflect generative models of the expected input via top-down pathways (Mumford 1992).

Resting scans are usually conducted under conditions in which subjects lie in a dark tube while fixating a cross in an otherwise blank display, which would not seem a fertile context for a perceptual prior. However, some aspects of an appropriate prior may not heavily depend on context. For example, recent work in monkey inferotemporal cortex indicates that individual faces can be coded by face cell assemblies whose firing rate is distributed along a small number of orthogonal dimensions (Chang & Tsao 2017). Therefore it is possible that the spontaneous activity patterns found in the present work reflect fluctuations along canonical low dimensional configurations.

### Synchronous fluctuations of representational content

The second important result is that multi-vertex patterns that putatively code a stimulus category fluctuate more synchronously at rest between visual cortical regions that prefer the same category (what we call pattern-based FC). Pattern-based FC between body-preferring regions was significantly larger when computed using a body-than scene-evoked pattern, while pattern-based FC between scene-preferring regions was significantly larger when computed using a scene-evoked than body-evoked pattern (Fig. 10).

An interesting result from the pattern-based FC analysis was that at rest different putative representational states, as indexed by multi-vertex patterns, fluctuated largely independently. Synchronous fluctuations associated with body-evoked patterns in body-preferring ROIs occurred independently of synchronous fluctuations associated with scene-evoked patterns in scene-preferring ROIs, as shown by the very low correlations between scene and body regions in the analysis of the Preferred Category FC matrix (Fig. 10). Therefore, resting activity across category-selective regions of visual cortex cannot be described in terms of a single representational state. These results provide new constraints on theories of the function of FC.

An interesting approach for identifying the representational content of resting FC has been reported in studies of early visual cortex, which have shown that resting FC respects the tuning of single voxels for polar angle, eccentricity, and low-level stimulus features (Arcaro et al 2015, Heinzle et al 2011, Raemaekers et al 2014, Ryu & Lee 2018). Most task-based studies of representation in higher-order visual and associative regions, however, have not involved measurements of voxelwise tuning functions but instead have identified task-evoked representations through measurements of regional patterns. Therefore, pattern-based FC (Anzellotti & Coutanche 2018, Chen et al 2018, Coutanche & Thompson-Schill 2013) can provide insights into the putative representational FC of spontaneous activity in high-level brain regions that are complementary to those provided by approaches based on the tuning properties of single voxels.

We suggest two additional ways in which pattern-based FC might inform studies of resting-state organization. First, pattern-based FC may help fractionate existing resting-state networks and identify the functional factors associated with that fractionation. For example, pattern-based resting FC between regions that prefer a particular category might depend on selectivity for features within the category, such as gender for face-preferring regions.

Second, pattern-based FC might uncover resting FC organizations that differ substantially from the normative whole-brain structure that has been described over the past decade (Cole et al 2016, Gordon et al 2016, Power et al 2011, Yeo et al 2011), although this structure does vary over individuals (Gordon et al 2017, Gratton et al 2018, Laumann et al 2015). In the current work, pattern-based FC was measured within category-preferring regions. Because regions that co-activate tend to show greater resting FC (Smith et al 2009), and because previous studies have shown that regions preferring the same category show preferential FC (Hutchison et al 2014, Stevens et al 2017, Turk-Browne et al 2010, Zhang et al 2009, Zhu et al), novel FC organizations were not expected. Accordingly, pattern-based FC matrices were moderately-to-strongly correlated with inter-regional vertex-averaged FC matrices. However, divergent FC organizations may be more likely in studies that use task-evoked activity patterns based on frequently occurring processes that combine different domains: for example, activity patterns based on integration of voice and face information during person-to-person interactions, visuomotor coordination during object manipulation, or biologically significant stimulus-reward or response-reward contingencies. Cross-domain pattern-based FC that cuts across standard networks might reflect synergies (Leo et al 2016, Santello et al 2013, Schieber & Santello 2004) or routines for implementing frequently occurring processes.

### Pattern-based FC and correspondence of resting and evoked activity patterns

We suggest that the synchronous fluctuations of representational content evidenced by pattern-based FC (Fig. 10) is partly responsible for the correspondence of stimulus-evoked activity patterns and spontaneous activity patterns that was observed in joint-ROIs (Figs. 7 and 8). The largest positive or negative similarity values for a resting frame in a joint-ROI will occur when the similarity values in the constituent ROIs on that frame are simultaneously large and have the same sign. Otherwise, across constituent ROIs the similarity values will tend to cancel or average to a lower value. Therefore, a larger spread of similarity values in a joint-ROI is more likely to be observed if the similarity values in the constituent ROIs fluctuate in a temporally correlated or synchronous fashion.

This mechanism explains the pattern of U90 values across the different joint-ROIs shown in Figure 7C. U90 values averaged across categories were highest in the face joint-ROI, intermediate for the body joint-ROI, and lowest for the scene joint-ROI, i.e. were inversely related to the number of constituent regions in each joint-ROI (2 for face, 5 for body, and 9 for scenes). The greater the number of constituent ROIs in a joint-ROI, the more the overall U90 value for the joint-ROI was decreased by sub-optimal synchronicity. As noted in the results section, in a two-factor ANOVA on U90 values with joint-ROI (body, face, scene) and Category (8 levels) as factors, the main effect of joint-ROI was significant. We additionally computed a single U90 value for each joint-ROI for each participant by averaging over categories. Paired t-tests on these averaged U90 values, with a Bonferroni-Holm correction for multiple comparisons (3 tests), indicated significantly larger U90 values within the Face than Body joint-ROIs (p<.0001), Face than Scene joint-ROIs (p<.0001), and Body than Scene joint-ROIs (p=.01).

In addition, since the multi-vertex activity patterns for a non-preferred category do not fluctuate as synchronously across constituent ROIs as those for a preferred category (Fig. 12), the resulting similarity values across frames in the joint-ROI for that non-preferred category will show less variation from zero, resulting in smaller U90 values. Therefore, within a joint-ROI, differences in U90 values between categories should be more reliable for joint-ROIs comprised of more constituent ROIs. This suggestion is also consistent with the results in Figure 7C, with the fewest significant differences found for the face joint-ROI and the most for the scene joint-ROI. Although other factors besides the number of constituent ROIs are clearly important in determining the category selectivity of U90 values in a joint-ROI, the larger point is that the greater pattern-based FC for the preferred category of the constituent ROIs increases the incidence of extreme similarity values between category-evoked and spontaneous patterns in a category specific fashion.

### Low- and high-level visual correspondences at rest

The spread of similarity values between resting activity patterns and stimulus-evoked patterns was determined by how well a stimulus activated the region, irrespective of whether the stimulus was more or less ecological. In many higher-level visual ROIs, stimulus preferences favored a particular whole-stimulus category (e.g. bodies) over another whole-stimulus category (e.g. scenes) or over the phase-scrambled category. Conversely, in early visual cortex, preferences favored stimuli that weighted low-level features, resulting in larger U90 values for scrambled than whole-stimulus categories. The larger U90 values for scrambled stimuli in early visual cortex do not contradict an overall framework in which resting activity patterns reflect the statistical distribution of features in the environment. Rather, this result suggests that resting activity patterns in regions that primarily extract low-level visual features are relatively independent of the patterns associated with higher-order features/statistics that define categories of more ecological stimuli.

Grid-scrambled objects showed greater U90 values than whole-objects in the Phase-scrambled joint-ROI and equivalent U90 values to whole-objects in the Whole-object joint-ROI (Fig. 8). This latter equivalence may have reflected the fact that the union of different category-preferential regions in the Whole-object joint-ROI eliminated or reduced the importance of features selective for a specific ecological category. Instead, the U90 value reflected features common to different ecological categories that were also present in grid-scrambled objects.

Therefore, the relationship between resting patterns and stimulus-evoked patterns can be driven by a variety of stimulus features that reflect local (e.g. contour-related features) or global (e.g. faces) stimulus characteristics depending on the tested regions.

### Limitations

Stimuli were not controlled for low-level variables that might have differentially activated visual regions. As noted, grid-scrambled stimuli may have included contour terminators to a larger degree than many whole-object stimuli, increasing the activation of early visual cortex. However, this factor was not explicitly controlled or manipulated. Also, stimuli were presented in a non-naturalistic context. Wilf et al. (Wilf et al 2017) have shown that in early visual cortex, resting FC patterns are better accounted for by movies than by standard retinotopic stimuli, while Strappini et al. (Strappini et al 2018) have shown that in higher-level visual cortex, resting FC patterns are better accounted for by movies than by static pictures of stimuli similar to those used here. Therefore, the present results may have underestimated correspondences between resting and evoked multi-vertex patterns.

## Acknowledgements

This work was supported by the National Institutes of Health RO1 MH096482 to MC.

